# Location Invariant Animal Recognition Using Mixed Source Datasets and Deep Learning

**DOI:** 10.1101/2020.05.13.094896

**Authors:** Andrew Shepley, Greg Falzon, Paul Meek, Paul Kwan

## Abstract

1. A time-consuming challenge faced by camera trap practitioners all over the world is the extraction of meaningful data from images to inform ecological management. The primary methods of image processing used by practitioners includes manual analysis and citizen science. An increasingly popular alternative is automated image classification software. However, most automated solutions are not sufficiently robust to be deployed on a large scale. Key challenges include limited access to images for each species and lack of location invariance when transferring models between sites. This prevents optimal use of ecological data and results in significant expenditure of time and resources to annotate and retrain deep learning models.
2. In this study, we aimed to (a) assess the value of publicly available non-iconic FlickR images in the training of deep learning models for camera trap object detection, (b) develop an out-of-the-box location invariant automated camera trap image processing solution for ecologist using deep transfer learning and (c) explore the use of small subsets of camera trap images in optimisation of a FlickR trained deep learning model for high precision ecological object detection.
3. We collected and annotated a dataset of images of “pigs” (*Sus scrofa* and *Phacochoerus africanus)* from the consumer image sharing website FlickR. These images were used to achieve transfer learning using a RetinaNet model in the task of object detection. We compared the performance of this model to the performance of models trained on combinations of camera trap images obtained from five different projects, each characterised by 5 different geographical regions. Furthermore, we explored optimisation of the FlickR model via infusion of small subsets of camera trap images to increase robustness in difficult images.
4. In most cases, the mean Average Precision (mAP) of the FlickR trained model when tested on out of sample camera trap sites (67.21-91.92%) was significantly higher than the mAP achieved by models trained on only one geographical location (4.42-90.8%) and rivalled the mAP of models trained on mixed camera trap datasets (68.96-92.75%). The infusion of camera trap images into the FlickR training further improved AP by 5.10-22.32% to 83.60-97.02%.
5. Ecology researchers can use FlickR images in the training of automated deep learning solutions for camera trap image processing to significantly reduce time and resource expenditure by allowing the development of location invariant, highly robust out-of-the-box solutions. This would allow AI technologies to be deployed on a large scale in ecological applications.

## 1. Introduction

Reducing the processing time of camera trap image analysis is fast becoming one of the most important drivers in the integration of computer science technologies and ecological research (Meek, Fleming et al. 2014, Fegraus and MacCarthy 2016, Meek, Ballard et al. 2019). Due to the increasing adoption of camera traps as a core survey method, the quantity of data collected by ecological practitioners has increased beyond that of any historical fauna survey method. These images are usually manually processed and analysed by practitioners or in some large-scale projects, by citizen scientists who classify them according to the goals of the study for which they were collected. Camera trap data can provide invaluable insight into a plethora of ecological information including species occurrence, activity patterns and behaviour (O’Connell, Nichols et al. 2011). While the benefits of this research represent a quantum shift in science, practitioners are restricted by the sheer volume of the data collected due to the substantial commitment of resources required to process it. This has prompted increasing research into the adoption of automated data-driven deep learning technologies for use in the task of image classification and analysis (Falzon, Meek et al. 2014).

Consequently, powerful, practical software tools such as ClassifyMe (Falzon, Lawson et al. 2020) have been developed, allowing ecology researchers to access and use cutting edge deep learning image classification solutions. However, such endeavours have not come without problems. Fully annotated ecological image data is sparse and not often readily or publicly available, which is severely limiting the general adoption of Artificial Intelligence in camera trap research and applications (Schneider, Taylor et al. 2018). Insufficient annotated data from a broad range of locations and species makes it very difficult for researchers to develop effective models capable of location and environmental invariance, resulting in the costly and inefficient task of training and developing models specific to each study (Falzon, Lawson et al. 2020). The lack of location invariance of modern detectors reduces their portability to camera trap sites not included in the training data, as they are highly susceptible to learning environmental factors such as camera angle, lighting, and vegetation as demonstrated by (Miao, Gaynor et al. 2019).

Thus, the aims of this research are twofold: (i) to evaluate publicly available sources of animal images such as FlickR to overcome barriers to image access and, (ii) to develop location invariant deep learning detector training strategies which allow the implementation of species recognition models across projects and regions. We will further propose optimisation strategies to achieve high precision results in a camera trap setting. This will allow a wealth of information usually lost due to the prohibitively large quantities of data to be gathered with ease, enabling more effective ecological management.

### 1.1 Manual Analysis and Citizen Science

The majority of camera trap image processing is achieved by manual analysis conducted by practitioners. Extremely large quantities of image data are collected by researchers world-wide and to-date every image sequence is interrogated by research teams, images uniquely renamed, and species tagged for analysis. Despite some camera trap systems being developed by users (see (Falzon, Lawson et al. 2020) for a contemporary summary), none have achieved high accuracy species recognition using automation software. Moreover, none have overcome the constraints of species-site based recognition due to the lack of location invariance of models trained specifically for each site and species cohort. This time-consuming task is alternatively achieved using citizen science. Essentially, a citizen scientist is a volunteer who contributes to scientific enquiry by collecting or processing image data (Nguyen, Maclagan et al. 2017). Large citizen science-based programs such as Zooniverse (www.zooniverse.org) enable the effective classification of millions of camera trap images (Jones, Allen et al. 2018). Citizen science projects have many benefits for researchers including customisation of projects and annotation requirements in accordance with the aims of their projects. However, the effectiveness of citizen science in meeting the data processing requirements of modern ecology is limited (Meek and Zimmerman 2016). The substantial investment of time and resources impedes timely image processing, causing delay between the data collection and interpretation stages, which may be detrimental to ecological management (Fox, Bourn et al. 2019). Furthermore, the need to upload large amounts of data onto publicly accessible websites may pose privacy risks (Sagarra, Gutiérrez-Roig et al. 2015) or poaching concerns and undermine the protection of rare or endangered species by revealing their geographical location and behavioural habits to poachers (Falzon, Lawson et al. 2020).

### 1.2 Deep Learning

Thus there is a clear need to automate the process of image annotation and classification in the context of camera trap images (Meek, Fleming et al. 2014, Meek, Ballard et al. 2015, Fegraus and MacCarthy 2016, Young, Rode-Margono et al. 2018). Computer vision technologies relying on deep learning, such as Deep Convolutional Neural Networks (DCNNs) have been at the centre of recent interest by researchers aiming to address the shortcomings of citizen science (Gomez Villa, Salazar et al. 2016, Norouzzadeh, Nguyen et al. 2017, Willi, Pitman et al. 2018). DCNNs are deep learning models trained on large datasets (thousands to millions) of images of interest until they can correctly classify images or detect objects within images (Zhao, Zheng et al. 2019). DCNNs are essentially learned algorithms which extract features from images and classifies them as belonging to a given class. Handcrafted features specified by researchers are not used, instead the features are ‘learned’ via updating of weights during training. When the DCNN is confident in the presence of an object in an image, it maps bounding boxes, segmentation masks, or classification labels to the image or object (Ren, He et al. 2015).

If a DCNN is very deep, consisting of many layers, it will have many trainable parameters (usually millions) which gives rise to the need for large annotated image datasets used in training these parameters from scratch. This is necessary for the network to learn complex features (Samala, Chan et al. 2016). Although DCNNs can be used to classify data with high accuracy, their usability can be limited by insufficient training data which may lead to overfitting, and consequently, inability of the model to generalise to new data (Zhao 2017). Another limiting factor is incorrect training processes, which may lead to the network learning environmental factors such as background and associating these features with a particular species (Miao, Gaynor et al. 2019). For example, a model trained on images of Reedbucks which have similar backgrounds may learn a camera-background/species correlation, resulting in misclassification and low recall (Miao, Gaynor et al. 2019). Thus, the intra-dataset variety in image background is a very important consideration when training models to be used across several traps.

In many fields, including security applications such as face recognition, DCNNs have achieved widespread success (Kouda, Morimoto et al. 2011, Ling, Jiyang et al. 2019) and have been broadly adopted by industry (Sánchez del Río, Conde et al. 2015). However, the adoption of DCNNs in ecological applications has been hindered by insufficient annotated data (Christin, Hervet et al. 2019), particularly rare species (Hao, Yang et al. 2019), and large variations between camera trap site geography which limits the useability of a network trained on data from one location, at another location (Willi, Pitman et al. 2018). Often, trained models learn features specific to a location or variation of a species, inhibiting their ability to generalise to other locations and species variations. Addressing this problem usually requires training of new models for each location or site (Everingham, Van Gool et al. 2010, Willi, Pitman et al. 2018, Falzon, Lawson et al. 2020), which is a time and resource intensive process. Thus, a generalised model which is able to achieve acceptable results in any given location for a particular species is an attractive option, particularly if it can be optimised by fine-tuning on a limited number of camera trap images.

### 1.3 Automatic Species Identification and Localisation

Early attempts at automated camera trap classification and object detection tasks using DCNNs were dependent on significant amounts of pre-processing (Yu, Jiangping et al. 2013) and resulted in relatively poor accuracy (Swinnen, Reijniers et al. 2014, Chen, Han et al. 2015). However, most modern solutions use minimal pre-processing, or automate pre-processing (Giraldo-Zuluaga, Salazar, Gomez, & Diaz-Pulido, 2017). Accuracy and recall attained by DCNN solutions is also increasing significantly, as large annotated datasets become available and progress is achieved in training methods, such as the adoption of transfer learning (Gomez Villa, Salazar et al. 2016, Willi, Pitman et al. 2018).

Transfer learning involves the repurposing of learned features for another task (Yosinski, Clune et al. 2014). This allows general features learned on a large, highly varied dataset such as ImageNet (Deng, Dong et al. 2009) which contains 3.2 million images, or Snapshot Serengeti (Swanson, Kosmala et al. 2015), which contains 7.3 million images to be transferred to a smaller, similar dataset containing only hundreds to thousands of images. Transfer learning has been shown to improve accuracy and the ability to generalise as well as reducing training time and the quantity of data needed (Khan, Hon et al. 2019). Its effectiveness in ecological camera trap applications has been established by (Norouzzadeh, Nguyen et al. 2017) and (Willi, Pitman et al. 2018).

### 1.4 Image Classification

The majority of camera trap image processing solutions achieve image classification rather than object detection (Gomez Villa, Salazar et al. 2016, Nguyen, Maclagan et al. 2017, Norouzzadeh, Nguyen et al. 2017, Willi, Pitman et al. 2018, Miao, Gaynor et al. 2019, Tabak, Norouzzadeh et al. 2019). Image classification is a process by which a whole image is labelled as containing a given object, for example, if a pig is featured in an image, the image will be labelled ‘pig’. However, image classification is limited in situations where an image contains more than one species, e.g. a pig and a wildebeest (Schneider, Taylor et al. 2018). Object localisation and object counting is also not effectively achieved by image classification, and models tend to struggle to distinguish between an empty frame, and a small background object (Yousif, Yuan et al. 2019). For this reason, this study focuses on object detection rather than image classification, as discussed in Section 1.5. For an overview of image classification methods, refer to Appendix S1.

### 1.5 Object Detection

In comparison to image classification, object detection is more useful because it allows more specific information to be extracted from images. Object detection is the process of locating and identifying one or more objects in an image. The model draws bounding boxes of varying classification confidence, together with an associated class label, around each object. It has many benefits, including object counting, and collection of data on specific individuals, which allows the study of reproduction, distribution, quantification, and comparison of behaviour across individual animals within a species group based on factors such as age and gender (Schneider, Taylor et al. 2018). Due to the substantial expenditure of time and resources necessary to annotate images with bounding boxes and class labels for training purposes, most deep learning solutions designed for camera trap image analysis conduct image classification rather than object detection. Like image classification models, object detection solutions either fail to address location invariance, or suffer the same dependency on familiarised camera trap dataset.

(Yousif, Yuan et al. 2019) employed sequence-level background subtraction using handcrafted Histogram of Oriented Gradient (HOG) (Dalal and Triggs 2005) features to localise moving objects in camera trap images. This study did not aim to identify animal species, instead simply distinguished between humans and animals, and eliminated empty frames. Although it achieved high accuracy in this task, its application was not extended beyond eastern North America. In contrast, our study provides a method by which camera trap images can be efficiently processed using a standardised DCNN procedure, removing the need for pre-processing, and successfully classifying animal species regardless of environmental factors and location.

A unique ecological image processing software solution for use on a laptop by field ecologists and wildlife managers was developed by (Falzon, Lawson et al. 2020). It provides object detection and localisation as well as species classification and object counting capabilities via training of YOLOv2 DarkNet-19 Deep Convolutional Neural Networks (DCNN) on both daytime and infrared imagery. It boasts fast processing speeds and acceptable accuracy, achieved on a local machine, within a dedicated on-demand application. Tailored models can be applied to trap sites in Australia, New Zealand, North America, Serengeti and the USA. However, optimal performance is only achieved when models are trained and developed for a specific environment, camera trap imaging configuration and species cohort. Thus, it suffers from lack of location invariance and robustness, as its accuracy and recall decrease when it is used outside the scope of the environments on which it was trained.

(Schneider, Taylor et al. 2018) addressed the problem of object detection in camera trap images, with the aim of identifying, quantifying and localising animal species. They used transfer learning to train a YOLOv2 (Redmon and Farhadi 2016) model, achieving recall of 93% and accuracy of 80.4% on the Reconyx (www.reconyx.com) and Snapshot Serengeti datasets. They also trained a Faster R-CNN model (Ren, He et al. 2015) achieving 76.7% recall and 72.2% accuracy. Like us, they used a pretrained COCO model to initialise transfer learning, using a small dataset; the Reconyx dataset contained 946 images of 20 species, while the Snapshot Serengeti dataset contained 4097 images of 48 species. However, the robustness of the model was not evaluated on out of sample images. It also suffered from class imbalance with lower accuracy and recall for classes with fewer instances. Our research indicates this limitation can be overcome by sourcing publicly available images from the internet.

### 1.6 Goals of this Study

i. To assess the value of non-camera trap domain imagery, in particular the publicly available photo-sharing website FlickR (https://www.flickr.com/) images in the training of deep learning models for camera trap object detection.
ii. To develop an out-of-the-box location invariant automated camera trap image processing solution for ecologists using transfer learning.
iii. To explore camera trap image infusion, a process that we proposed and coined in this work to refer to the use of small subsets of camera trap images in the optimisation of FlickR trained deep learning models for high precision ecological object detection.

## 2. Datasets and Methodologies

### 2.1 Datasets

The context of this study focuses on the detection of Suidae (pigs, hogs and boars) using single-class object detectors. Monitoring of Suidae is of great importance due to the significant role they play across global ecosystems and their host status for a range of diseases such as Swine Fever, which are major threats to agricultural industries.

This study used a dataset of images collected from FlickR. FlickR was chosen due to the ease at which large numbers of images could be downloaded using the FlickR API, and its established status as a reliable source of images for deep learning (Everingham, Van Gool et al. 2010, Lin, Maire et al. 2014). It also used five camera trap datasets containing images of variations in species of pig and warthog, collected by different research teams in Australia, Europe, North America, South Africa and Tanzania. These images were provided without bounding box annotations, which resulted in the need to develop a consistent annotation protocol for training and testing of the DCNN models. Negative samples were explicitly added to the training sets to allow the network to better discriminate between foreground features belonging to the Suidae class, and those belonging to out of sample species. For more information on the negative sampling dataset and procedure used, refer to Appendix S2.

#### i. FlickR

As previously mentioned, one of the biggest challenges faced by ecological researchers when deploying DCNNs for camera trap image processing is the lack of location and environmental invariance of trained models. As early as 2008, studies in contextual object detection (Hoiem, Efros et al. 2008, Sudderth, Torralba et al. 2008) examined the consequences of ‘unintentional regularities’ in datasets resulting in object detectors learning associations between objects and their background, inhibiting their ability to detect objects out of context. (Everingham, Van Gool et al. 2010) noted that classifiers tend to learn the context of an object rather than model the appearance of the object. Thus, when the object is dissociated with its context, e.g. pigs in a location or environment not included in the training set, the classifier fails to detect them as they are out of their context. They established that most classification methods make extensive use of image composition and context, with performance dropping significantly when objects are out of context. This research has been validated in an ecological context by (Miao, Gaynor et al. 2019) which observed that networks have a tendency to learn background features as elements of an object if image background and context is not highly varied. It is therefore essential to broaden the context of animal imagery to extend beyond a restricted range of camera traps to ensure robustness, and location and context invariance.

This phenomena of contextual association was investigated by (Everingham, Van Gool et al. 2010) who observed that images taken by researchers for a specific purpose, such as the collection of camera trap images by ecologists, creates an inner dataset bias. This results in the development of models incapable of generalisation to other camera trap contexts. On this basis, we postulate that collection of camera trap images for DCNN training mimics collection of images under laboratory or controlled conditions – aspects such as lighting, camera angle, distance of animals from the camera, and background features are consistent across many images in the training set, which would encourage contextual association with species identification. This is supported by (Willi, Pitman et al. 2018) who noted that their models, trained on camera trap images, would need to be retrained for use in other camera traps which did not form part of the training set.

One method used by researchers to overcome this challenge is the use of public photo sharing websites such as FlickR, Pinterest (www.pinterest.com), Imgur (www.imgur.com), 500px (www.web.500px.com) and Pixabay (www.pixabay.com) which showcase images exhibiting an extensive range of contextual features (Everingham, Van Gool et al. 2010, Lin, Maire et al. 2014). FlickR is used by amateur photographers to upload and share images with searchable metadata and keywords. Notably, FlickR contains a broad range of non-iconic images, highly suitable for DCNN training. Iconic images contain a single, large, unobstructed view of an object, usually positioned in the centre of the image (Lin, Maire et al. 2014). In contrast, non-iconic images feature large variations in background, occlusion, lighting, camera angle and context, as illustrated by Figure 1. This renders them invaluable in the development of DCNN object detectors capable of generalisation (Torralba and Efros 2011), thus making them very useful in real life object detection (Hoiem, Chodpathumwan et al. 2012). This accounts for their usage in extensively used object detection datasets and benchmarks such as MS COCO (Lin, Maire et al. 2014) and PASCAL VOC (Everingham, Van Gool et al. 2010).

**Figure 1:**
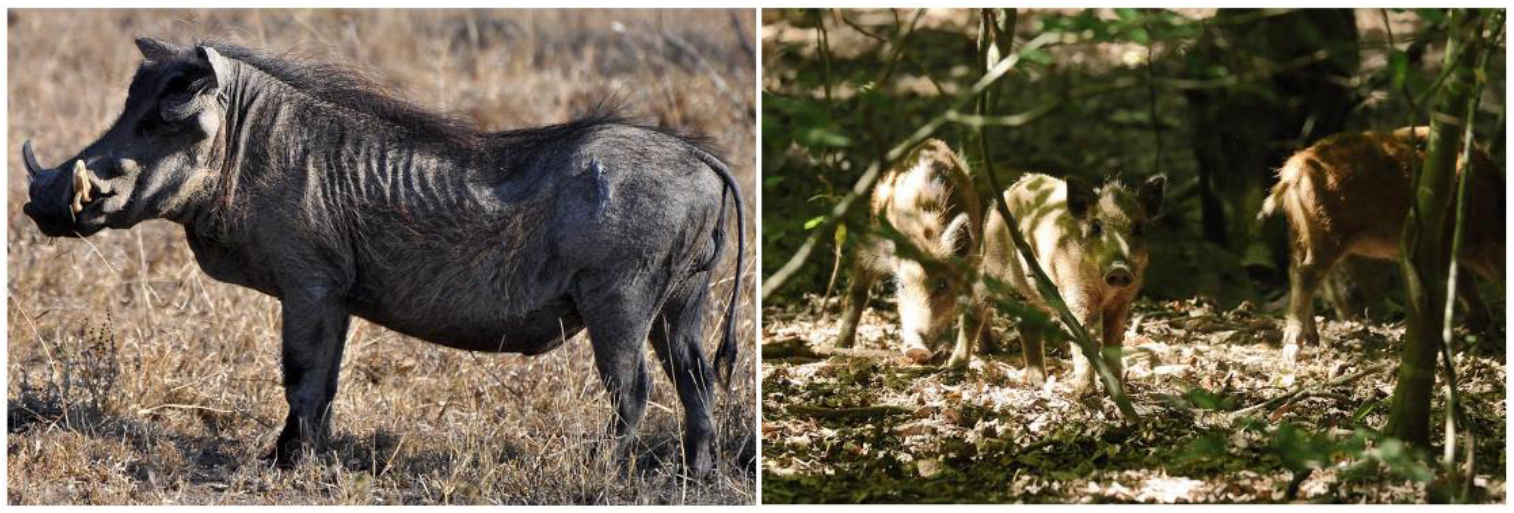
Left: an iconic image of a boar. Right: a non-iconic image of three pigs (sourced from FlickR).

Ecologists are tasked with a similar challenge – to classify and localise animals in a wide spectrum of natural images (Norouzzadeh, Nguyen et al. 2017, Willi, Pitman et al. 2018, Tabak, Norouzzadeh et al. 2019). To this end, we propose the general usage of FlickR images, rather than camera trap images, as the primary source of data for the task of DCNN model development for automated camera trap image analysis. We downloaded a dataset of approximately 700 images, restricting our search to Creative Commons licensed images. We noted however that many images, particularly infrared images, were protected by copyright, preventing their usage in this project.

As this research involved transfer learning which only requires in the order of hundreds to thousands of images (Khan, Hon et al. 2019), we considered the size of the dataset adequate for our purposes. After processing and annotation, our final dataset contained 606 images, including 1061 annotations of the animal family Suidae (pigs). Images were obtained from FlickR using the FlickR API. We used a set of keywords shown in Table 1, allowing for 200 images per search query. Specific and general association terms were used to ensure relevance of results and include sufficiently varied contextual images. The time taken to download the entire dataset was approximately 15 minutes.

**Table 1:** Search queries or keywords used to download images of pigs from FlickR.

| Search Queries |  |
| --- | --- |
| Australian+feral+pig | Warthog |
| Noxious+pest+pig | Sus+scrofa |
| Feral+pig+hunting | Wild+pig |
| Feral+pig+control | Wild+boar |
| Phacochoerus+africanus | Wild+hog |
| Feral+pig | Sanglier |
| Wild+pig |  |

The final dataset was compiled by manually removing all images which did not contain pigs. Duplicates and near duplicates were also removed. Near duplicates are images that have strong visual similarity, containing only small photometric distortions, variations and occlusion (Everingham, Van Gool et al. 2010). A maximum of 3 images taken by the same photographer during the same period of time were included, to maximise variability of images (Lin, Maire et al. 2014). A sample of the final dataset is shown in Figure 2.

**Figure 2:**
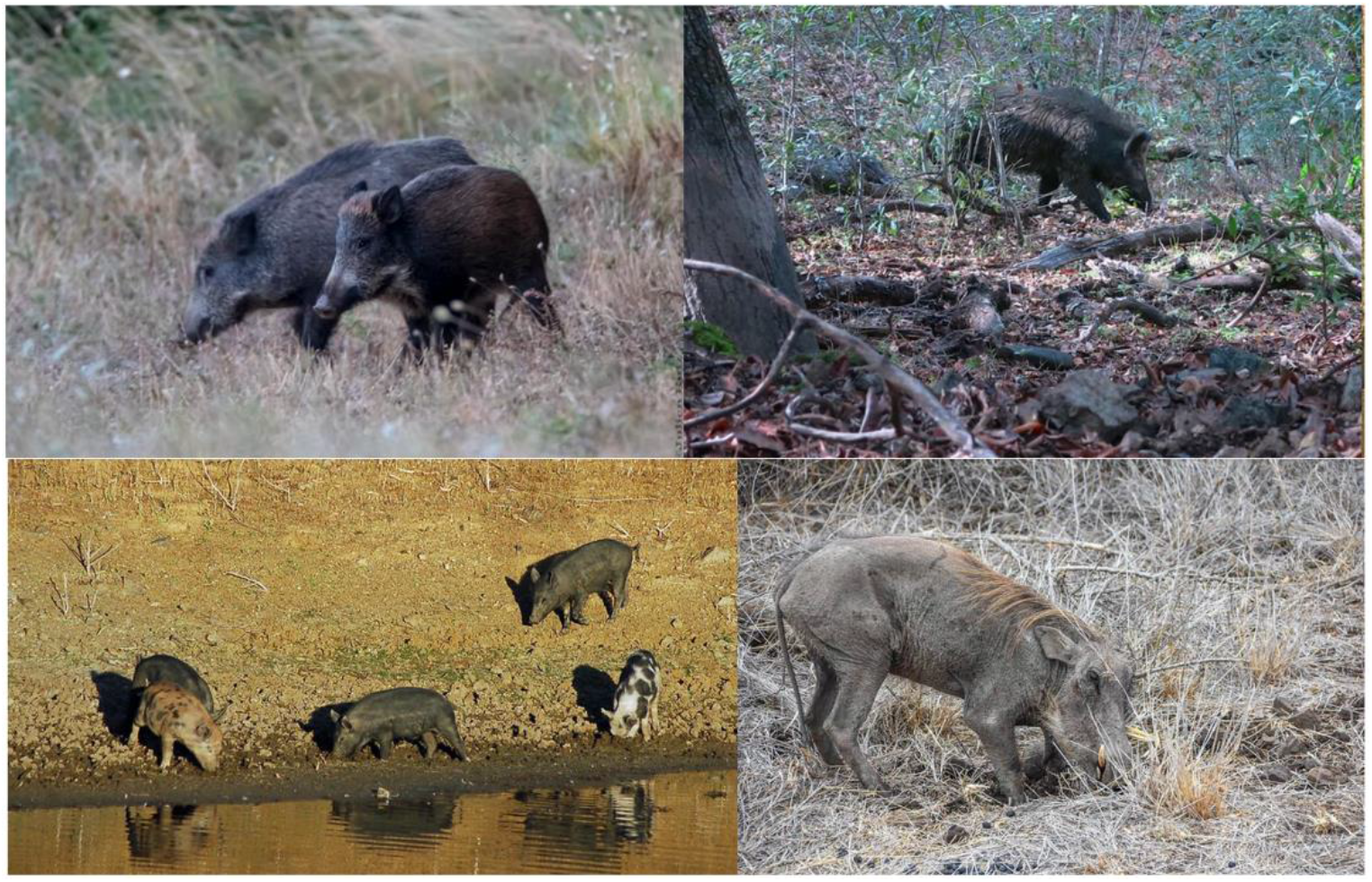
Examples of images obtained from FlickR for location invariance training in the task of Suidae detection.

Each image in the final dataset was annotated with bounding boxes and corresponding class labels. Bounding box annotation involves the positioning of an axis aligned box surrounding an object. We used the graphical annotation tool `labellmg` (Tzutalin 2015) for annotation, saving annotations in PASCAL VOC format. All bounding box annotations were completed by one researcher to improve consistency in bounding box placement, and to minimise errors.

#### ii. Camera Trap Datasets

The camera trap data used in this study was collected from a variety of groups across a spectrum of species, landscapes and habitats utilising a wide variety of camera models, configurations and image resolutions, as illustrated by Figure 3. This study therefore constitutes a wide sample of camera trap images representative of Suidae across projects. Most important to our study is the qualitative characteristics of the camera trap images obtained rather than the technical configurations.

**Figure 3:**
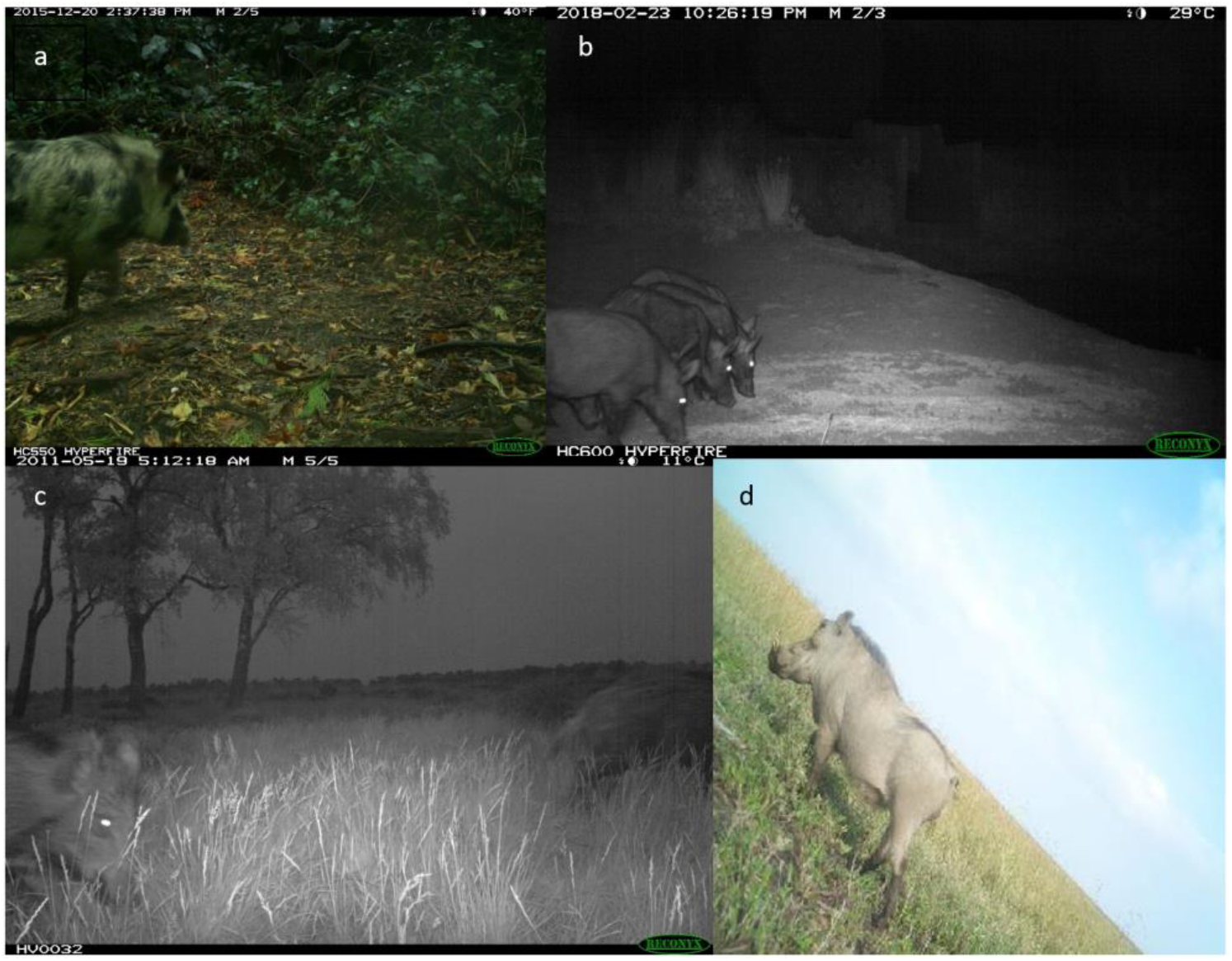
Examples of camera trap images from four out of the five locations used in this study; North America (a), Australia (b), Europe (c), and Tanzania (d).

The Australian dataset comprised of 6 locations in Northern NSW, Australia, obtained during feral pig trapping and control operations. The majority of images were captured at night (infrared) due to pig activity patterns. Some images had high occlusion with many pigs per image and pigs were often far from the camera.

The Tanzanian dataset was sourced from the *Snapshot Serengeti* project (Swanson et al. 2015). A specific subset of the original data was utilised from the Data Repository for the University of Minnesota https://doi.org/10.13020/D6T11K utilised by (Willi, Pitman et al. 2018) and released under a CC0 1.0 Universal Public Domain Dedication https://creativecommons.org/publicdomain/zero/1.0/.

The South African dataset was sourced from the Camera CATalogue project conducted by Panthera (www.panthera.org) in South Africa. This dataset was also provided from the Data Repository for the University of Minnesota https://doi.org/10.13020/D6T11K utilised by (Willi, Pitman et al. 2018) and released under a CC0 1.0 Universal Public Domain Dedication https://creativecommons.org/publicdomain/zero/1.0/. The ‘warthog’ class was extracted and utilised in this study. Note that the original camera trap images were resized to 330 × 330 pixels. Both the Tanzania and South Africa were semantically similar, containing mostly daytime images characterised by low occlusion, and few pigs per image.

The Europe dataset comprised of a dataset of 501 images of boars sourced from the Missouri Camera Traps dataset (Zhang, He et al. 2016). These images were very challenging, with highly cluttered and dynamic scenes. This dataset was released under the Community Data License Agreement (permissive variant). Although this dataset contained 25,000 images representing 20 species, only images of ‘wild boar’ were extracted for this study. Image spatial resolutions varied from 1920 x 1080 to 2048 x 1536. These images were not resized.

The North America dataset was sourced from the North American Camera Trap Images dataset (Tabak, Norouzzadeh et al. 2019). This dataset was of high quality, containing images from five locations across the United States. The entire dataset was made up of 3.7 million images of 28 classes. A subset of 514 images of wild boars was derived from this dataset for use in this project.

We randomly assigned images from each dataset to two subsets; a training set (90% of the data), and a validation set (10% of the data) as illustrated by Table 3. Models were trained on the training set. The validation split was used to monitor training to reduce overfitting. This was achieved by continuing training until accuracy stopped improving or began to decline. A test set for each dataset was not required as testing was only conducted on out of sample data.

**Table 3:** Distribution of dataset imagery across training and validation subsets.

|  | Dataset Sources | Number of Images | Training Split | N° of Training Instances | Validation Split | Test Locations |
| --- | --- | --- | --- | --- | --- | --- |
| Camera Traps | Australia | 589 | 530 | 1698 | 59 | North America, Europe, Tanzania, South Africa |
|  | North America | 514 | 463 | 779 | 51 | Australia, Europe, Tanzania, South Africa |
|  | Europe | 501 | 451 | 614 | 50 | Australia, North America, Tanzania, South Africa |
|  | Tanzania | 574 | 517 | 647 | 57 | Australia, North America, Europe, South Africa |
|  | South Africa | 559 | 503 | 601 | 56 | Australia, North America, Tanzania, Europe |
|  | FlickR | 606 | 546 | 961 | 61 | Australia, North America, Tanzania, South Africa, Europe |

### 2.2 Training Method

Details of the model architecture and training parameters are provided in Appendix S3. Additional information on transfer learning is also provided. The experiments outlined in this Section were verified on a multi-class application documented in Appendix S5. The multi-class application also involved training of a RetinaNet model on 4 classes (goat, pig, kangaroo and fox), followed by evaluation on a real-life camera trap dataset.

#### i. Model Transferability – Training on FlickR to achieve Location Invariance

The primary focus of this research was to develop deep learning models trained using non-camera trap imagery and evaluate their performance across diverse locations. Thus we conducted a comparison of the transferability of a model trained only on FlickR images, against five models trained on data from five separate locations, and five models trained on combinations of camera trap datasets from different countries. In this study, transferability refers to the ability of a trained model to be deployed on out of sample test data. Out of sample test data refers to animal imagery collected from trap locations and sites not included in the training data.

We trained five Keras-RetinaNet models (Lin, Goyal et al. 2018) on combinations of camera trap images from various camera traps and locations as outlined in Table 4. Each model was trained on a dataset made up of a combination of four camera trap datasets, supplemented with 788 negative samples (see Table 2). The training process for each model was monitored using a validation set made up of a combination of each corresponding validation set in Table 3. For example, CT Model 1 was trained on a combination of the training sets belonging to each of South Africa, Tanzania, Australia and Europe. It was validated on a dataset comprised on the validation sets for each of South Africa, Tanzania, Australia and Europe. Each model was then tested on an out of sample test set comprised of the entire set of images from the location the model was not trained on, for example, CT Model 1 was trained on South Africa, Europe, Tanzania and Australia datasets, and was tested on the North America dataset.

**Table 4:** Training, validation and testing profiles for each camera trap model. Note all training sets each include 788 negative samples

| **Camera Trap Models** | **Locations** | **N° of images** |  | **Out of Sample Test Set** |  |
| --- | --- | --- | --- | --- | --- |
|  | **Training/validation** | **Training** | **Validation** | **N° of images** | **Location** |
| *CT Model 1* | South Africa, Tanzania, Australia, Europe | 2789 | 222 | 514 | North America |
| *CT Model 2* | South Africa, Tanzania, Europe, North America | 2722 | 214 | 589 | Australia |
| *CT Model 3* | South Africa, Australia, Europe, North America | 2735 | 216 | 574 | Tanzania |
| *CT Model 4* | Australia, Europe, North America, Tanzania | 2649 | 217 | 559 | South Africa |
| *CT Model 5* | South Africa, North America, Australia, Tanzania, | 2801 | 223 | 501 | Europe |

The FlickR model was trained on 606 images, containing 961 instances of the class ‘pig’ (see Table 3). Like the camera trap models, the training set included 788 negative samples. No camera trap images were included in training or validation. The model was tested on each of the out of sample test sets identified in Table 4. These out of sample test sets were comprised of images, each containing at least one instance of the class ‘pig’.

Single location camera trap models were also trained to evaluate the transferability of models trained on data from one country or region to other countries or regions. For example, the transferability of a model trained solely on camera trap images sourced from sites in South Africa was evaluated on camera trap data with high dataset similarity, such as Tanzania, and dissimilar datasets such as Australia, Europe and North America. The training, validation and test information for the models is provided in Table 3. All training sets were supplemented with 788 negative samples.

#### ii. Optimisation of FlickR Training Using Camera Trap Image Infusion

We conducted further experiments to assess an optimization process that would allow ecologists to improve their FlickR model performance with a minimal infusion of camera trap images into the FlickR training set. The FlickR model (0% image infusion) was trained as described above. Four additional models were trained for each location, with incremental infusion of camera traps into training datasets to observe the impact of the introduction of trap images into the FlickR training. Subsets of images from each camera trap training set were added in increments of 5% as shown by Table 5.

**Table 5:** Incremental infusion of camera trap images into the FlickR training. The training set for each model included the original 606 images of ‘pig’ from the FlickR training set, as well as 788 negative samples.

|  | Model | Infusion (%) | N° infusion images | N° infusion instances | Training set size | Test set size |
| --- | --- | --- | --- | --- | --- | --- |
| Australia | CT Model 1 | 5 | 30 | 108 | 1424 | 410 |
|  | CT Model 2 | 10 | 60 | 197 | 1454 |  |
|  | CT Model 3 | 15 | 90 | 263 | 1484 |  |
|  | CT Model 4 | 20 | 120 | 359 | 1514 |  |
| South Africa | CT Model 1 | 5 | 30 | 34 | 1424 | 383 |
|  | CT Model 2 | 10 | 60 | 73 | 1454 |  |
|  | CT Model 3 | 15 | 90 | 103 | 1484 |  |
|  | CT Model 4 | 20 | 120 | 137 | 1514 |  |
| North America | CT Model 1 | 5 | 30 | 42 | 1424 | 343 |
|  | CT Model 2 | 10 | 60 | 79 | 1454 |  |
|  | CT Model 3 | 15 | 90 | 130 | 1484 |  |
|  | CT Model 4 | 20 | 120 | 171 | 1514 |  |
| Tanzania | CT Model 1 | 5 | 30 | 35 | 1424 | 397 |
|  | CT Model 2 | 10 | 60 | 66 | 1454 |  |
|  | CT Model 3 | 15 | 90 | 98 | 1484 |  |
|  | CT Model 4 | 20 | 120 | 131 | 1514 |  |
| Europe | CT Model 1 | 5 | 30 | 46 | 1424 | 331 |
|  | CT Model 2 | 10 | 60 | 85 | 1454 |  |
|  | CT Model 3 | 15 | 90 | 109 | 1484 |  |
|  | CT Model 4 | 20 | 120 | 142 | 1514 |  |

Each subset was added to the FlickR/Negative sample training set. Training was validated on a combination of the FlickR validation set and the camera trap validation set (see Table 3). Testing was conducted on the unused camera trap images. For example, the Australian camera trap training set was comprised of 530 images. The largest infusion percentile used was 20% of the size of the FlickR dataset, which means 120 images were used for infusion training. The remaining 410 images were used for testing.

Preliminary investigations using randomly selected subsets produced inconsistent results, largely due to the high number of partial images of pigs as shown by Figure 4, and the repetition of very similar images caused by the camera trap taking multiple images of the same event (capture event), as illustrated by Figure 5.

**Figure 4:**
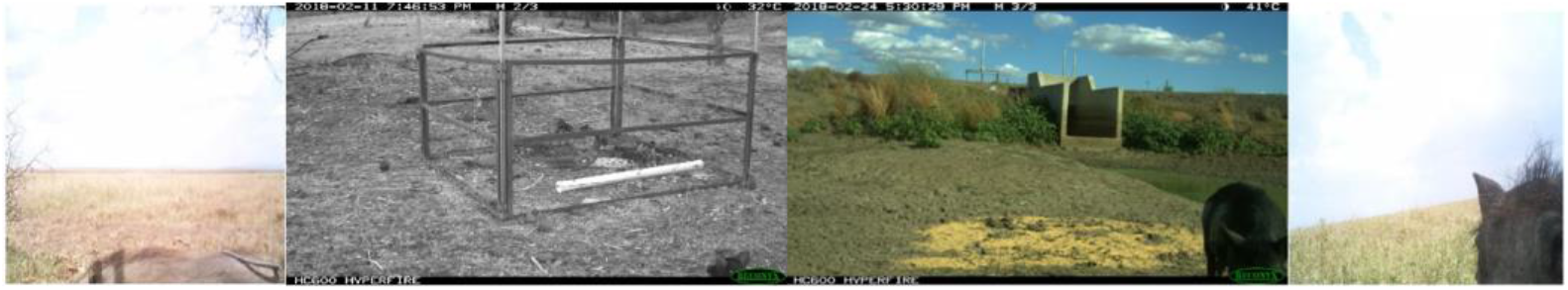
Inclusion of partial images in training cause the DCNN to overenthusiastically classify objects which resemble partials as pigs. Thus, they were removed from the training set (but retained in testing set).

**Figure 5:**
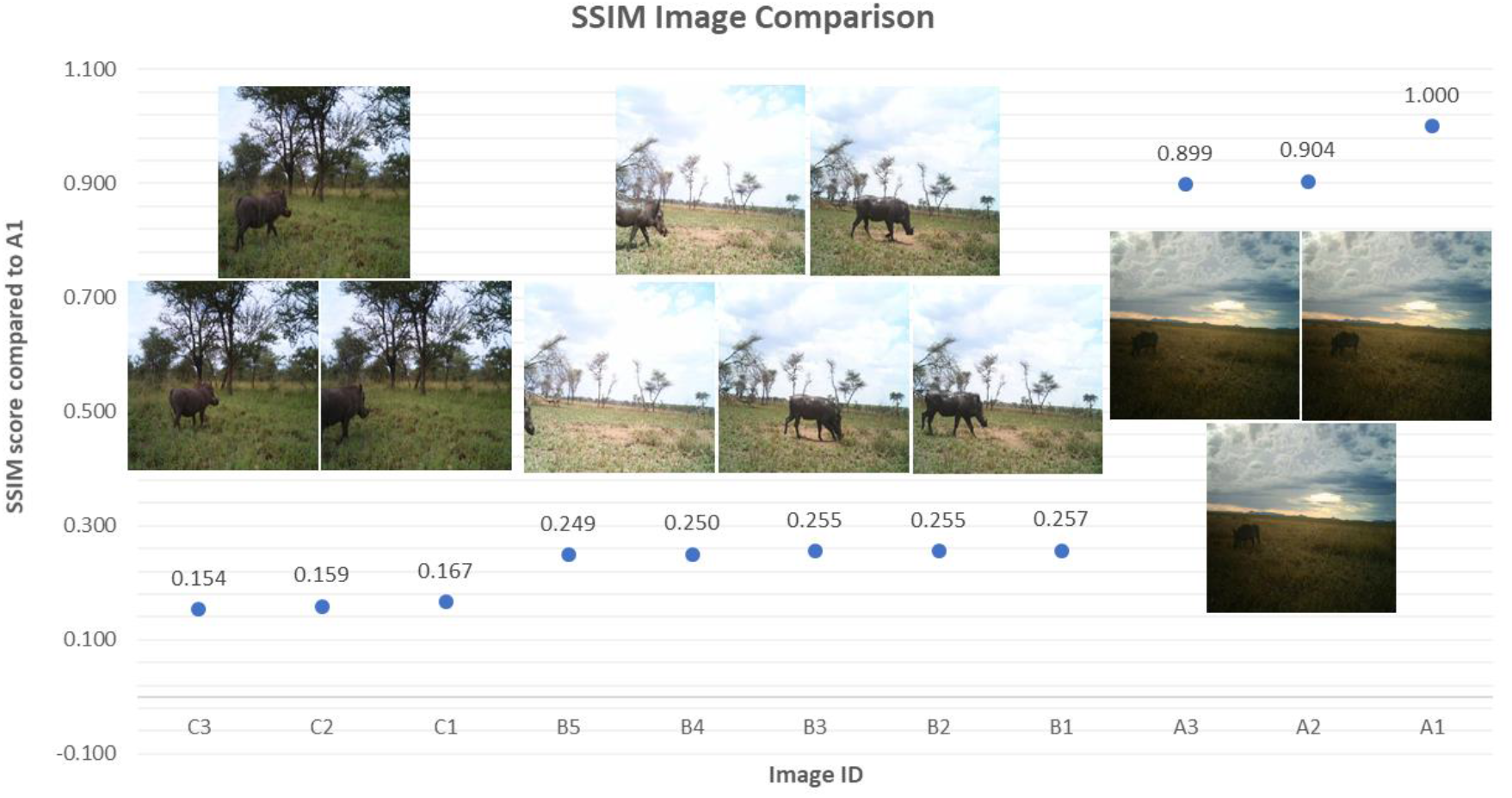
Graphical illustration of image clustering using an SSIM clustering algorithm

To address this, a Structural Similarity Index (SSIM) (Zhou, Bovik et al. 2004) clustering algorithm was developed to automatically sort images based on SSIM values. This algorithm is provided in Appendix S4. It allowed us to randomly select a frame from each cluster of images (usually one capture event, or different capture events with very similar properties). Our research indicates that image pairs with an SSIM value above 0.4 have sufficiently high similarity to be clustered. For example, Figure 5 illustrates the output of the SSIM algorithm graphically, clearly showing the three clusters formed by visually similar images. The image denoted by the arrow (the test images) is compared to each other image, with values closest to 1 indicating high similarity with the test image. This method allows researchers to compile highly varied datasets, improving learning, and minimising the need for extensive, time-consuming image sorting and annotation.

#### iii. Model Evaluation

To evaluate the performance of our models, Average Precision (AP) results will be provided. AP is calculated as documented in the PASCAL VOC benchmark (Everingham, Van Gool et al. 2010). A high AP indicates that the model is detecting the majority of objects at high accuracy. Accuracy is measured using Intersection over Union (IoU), which is a measure of the overlap between the detection box and the ground truth.

## 3. Results

### 3.1 Model Transferability – Training on FlickR to achieve Location Invariance

The results of the model transferability experiments are presented in Figure 6. The Flickr Model achieved mAP results between 67.21% – 91.92%, while the single location Camera Trap model performance ranged from 4.42-90.80%. The combined location camera trap models achieved mAP results ranging from 68.96-92.75% on out of sample camera trap test sites.

**Figure 6:**
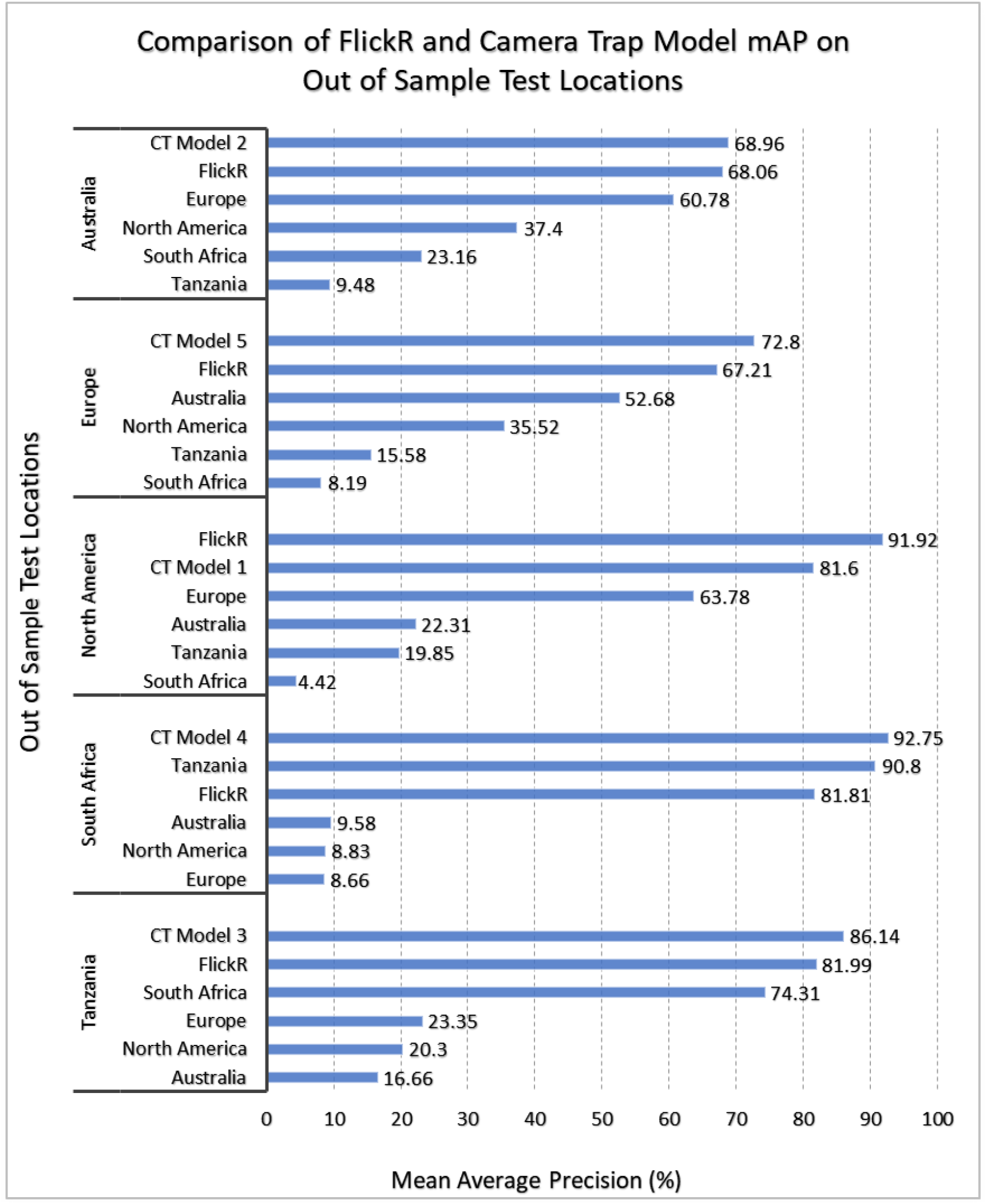
Comparison of APs achieved by models trained on single country/region camera trap datasets, mixed country/region datasets and a model trained on the FlickR dataset when tested on out of sample datasets.

CT Model 1 achieved a mAP of 81.60% on the North America test set (514 images of class ‘pig’). This may be explained by the high dissimilarity in background, lighting and species variation. As shown by Figure 7, images from the North America camera traps were highly varied and challenging, containing heavily wooded or snow-covered backgrounds, with dappled sunlight, high contrast, and the species *Sus scrofa.* In contrast, the camera trap sites from Australia, South Africa, Europe and Tanzania contain grassy or dirt backgrounds, night vision and more evenly distributed sunlight, and the species *Phacochoerus africanus* (South Africa and Tanzania) or *Sus scrofa* (Australia and Europe). As the FlickR training dataset contained several images of pigs in wooded and snow settings, and had a wide range of backgrounds, lighting and species variations, it was able to better generalize, achieving an AP of 91.92%. These results indicate that the FlickR trained model is more robust to a wider range of camera trap images.

**Figure 7.**
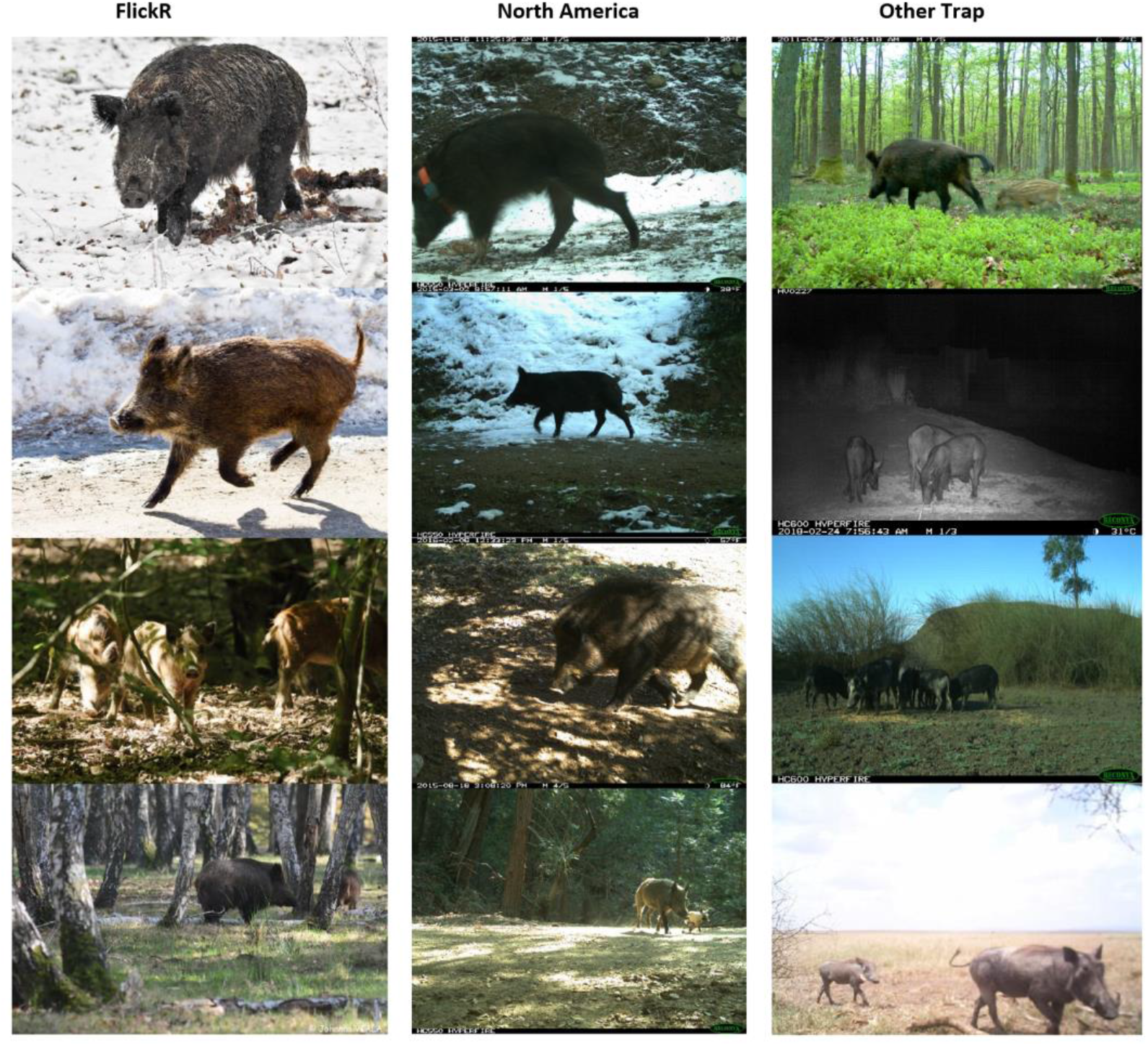
Similarities between some images in the FlickR dataset (left) and those contained in the North America test set (centre) allow it to better transfer to achieve a high mAP (91.92%) in comparison to the single location and mixed location camera trap datasets (right), which contain images which are very different to those in the North America test set.

CT Model 2 achieved a mAP of 68.96% when evaluated using the Australia dataset. The model trained on FlickR images achieved a similar mAP (68.09%). One factor which could explain the low mAP was the lack of infrared images in the FlickR training set, the latter of which formed the majority of Australian trap images. This result is nevertheless suboptimal but indicates that the FlickR model is more robust and location invariant when compared to the camera trap model, due to the fact that even without training on infrared images, it was able to achieve a very similar result to CT Model 2, which included infrared images (Europe and Missouri datasets). In the next section, we will present results of the optimisation experiments were conducted to investigate how the FlickR model can be improved with minimal infusion of Australian trap images.

Similarly, CT Model 5 obtained a mAP of 72.80% when tested on the Europe dataset. The FlickR trained model achieved a mAP of 67.21%. The Europe dataset featured mostly infrared images, putting the FlickR trained model at a disadvantage, compared to CT Model 5, which benefited from the inclusion of the very difficult Australian infrared images within its training set. The superior performance of CT Model 5 highlights the need for inclusion of both daytime and infrared images in camera trap training, which is one shortcoming when using FlickR as an image source – most infrared images on FlickR are restricted by copyright, limiting data availability to daytime or colour night-time images.

When tested on the Tanzania dataset, CT Model 3 achieved an AP of 86.14%, outperforming the FlickR trained model by 4.15%. This was due to the high similarity between warthogs in South Africa and Tanzania, and their corresponding visual backgrounds. This result suggests that camera trap models can achieve higher accuracies only when there is strong similarity between species and camera trap locations. This is evidenced by the similarly high performance of a model trained only on the South Africa dataset, on the Tanzania test set (74.31%). However, the FlickR trained model is generally far more robust on unseen datasets due to the significant variety in training images, which allows it to better generalize to unseen datasets.

CT Model 4 achieved an AP of 92.75% when assessed on the out of sample South Africa dataset. The FlickR Model achieved an AP of 81.81%. It is noteworthy that the superior performance of CT Model 4 is due to the abovementioned similarity between the South African and Tanzanian warthogs and their visually similar backgrounds. The model trained only on Tanzanian data achieved a mAP of 90.80% on the South Africa test set, while the models trained on data from other countries/regions achieved mAP results ranging from 8.66-9.58%, suggesting that the addition of the training data from Europe, North America, and Australia contributed very little to the high performance of CT Model 4.

### 3.2 Optimisation of FlickR Training Using Camera Trap Image Infusion

The previously mentioned results indicate that the FlickR trained model can be used to process images collected by any camera trap site, even if the FlickR training set does not contain any trap images, and large dissimilarities exist between the FlickR data and the camera trap site. For example, the FlickR training set did not include any infrared images. Despite this, the model trained on this data was able to detect many pigs in infrared images, as shown by Figure 8. Notably, the Australian dataset was very challenging, containing poor quality infrared images, which accounts for the relatively low AP achieved by the FlickR model on the Australian test set.

**Figure 8:**
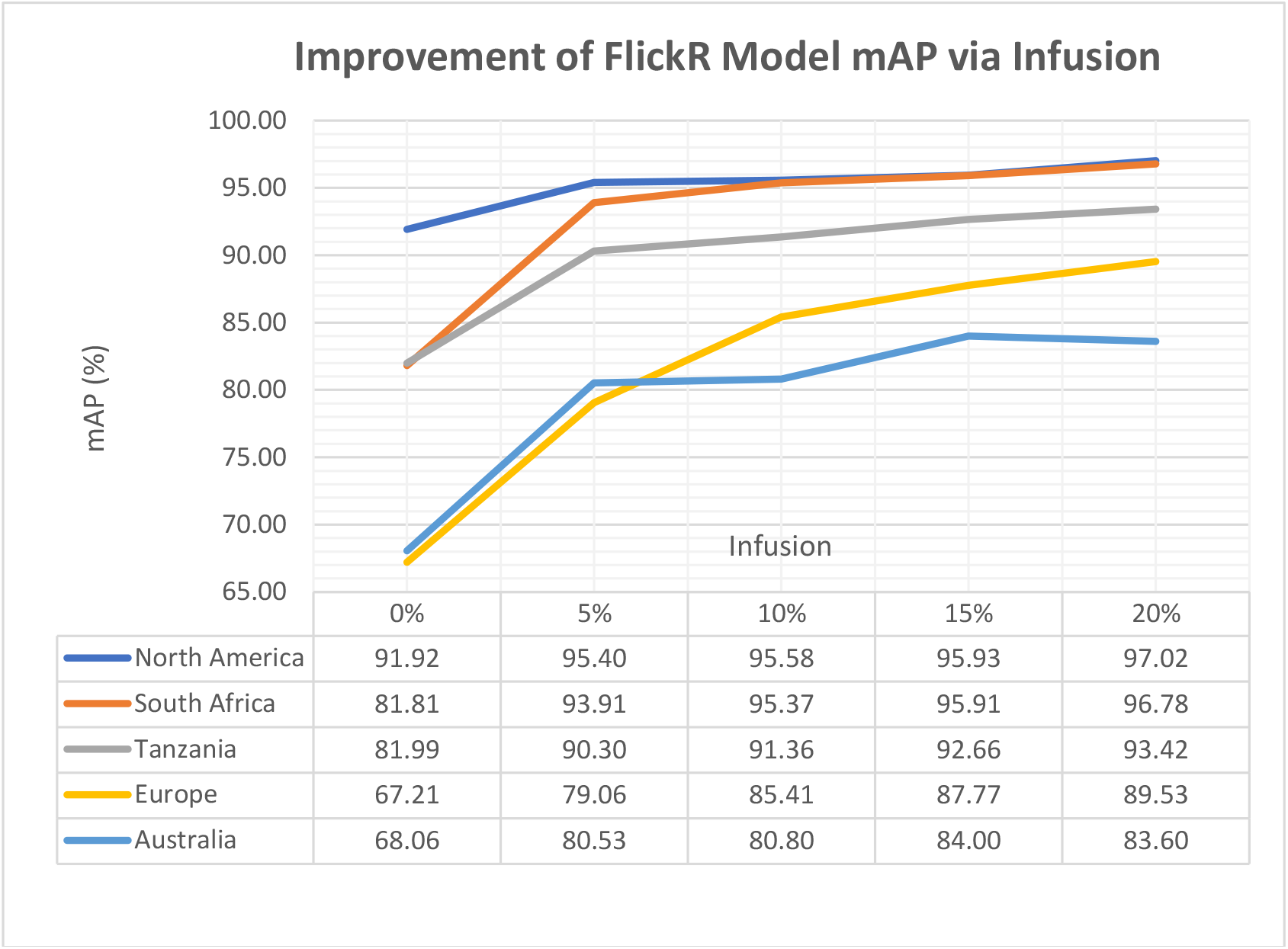
Impact of infusion of trap images into FlickR training on model AP.

It is clear however that a mAP of 68% is not sufficiently high for practical purposes. Thus, we present the results of our optimisation experiments (see Figure 8). These results indicate that the addition of a small percentage of camera trap images into the FlickR training set significantly improves performance. In all cases, the infusion of 5-10% trap images into the FlickR training resulted in a significant increase in mAP (3.66-18.20%). Increasing the infusion set size from 10% to 20% did improve mAP results in 4 out of 5 cases, however improvements were not substantial (1.41-4.12%).

Interestingly, the mAP of the Australian model began to decrease with the infusion of more than 15% trap images into training. We believe this is due to the poor image quality (see Figure 9) and lack of intra-dataset variability within the Australian image dataset. In contrast, the mAP of the other infusion models continued to increase, albeit slightly. Notably, the models which benefited the most from infusion, were those infused with camera trap imagery containing infrared images, and/or the species *Phacochoerus africanus*. This may be a consequence of the relatively small percentage of *Phacochoerus africanus* in the FlickR training set compared to *Sus scrofa.* This suggests that it is of significant importance to achieve uniformity in subspecies distribution in the dataset to achieve an optimal AP. Thus, class imbalance is not only detrimental across species (Willi, Pitman et al. 2018), but also appears to be within species variations. Another cause of the continued increase is the greater intra-dataset variability in the South Africa and Tanzania datasets – these sets contained images from more traps, with different backgrounds, camera angles, lighting etc. which made the addition of more images useful to the DCNN training set.

**Figure 9:**
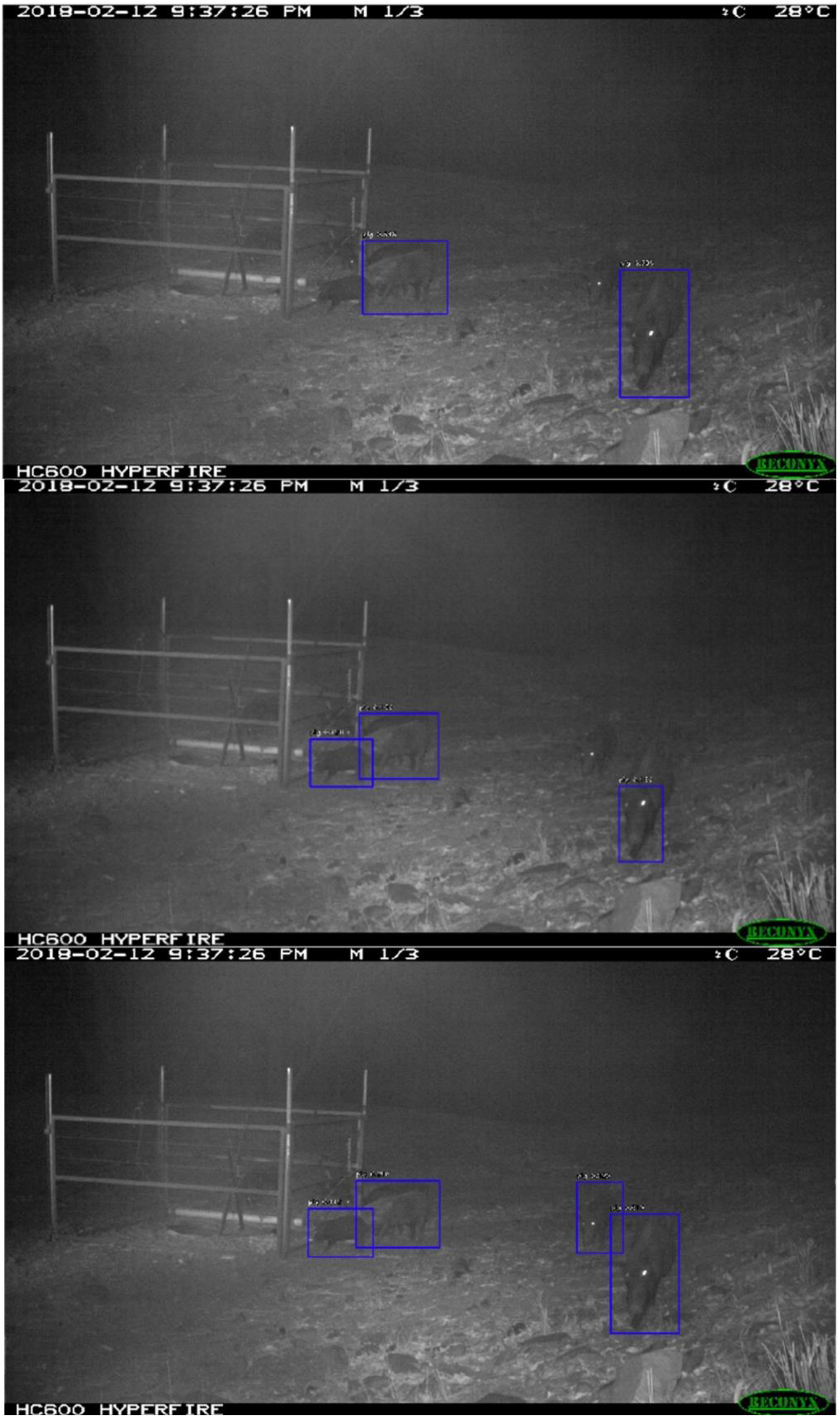
Top: output from the camera trap model on sample images from the Australian test set (trained on South Africa, Europe, North America and Tanzania). This model only detects 2 out of 5 pigs, whereas the FlickR Model detects 3 out of 5 pigs (centre). The 5% infusion training model detects all 4 clearly visible pigs (bottom), only missing one highly occluded pig (far right).

A qualitative example of the benefit of camera trap infusion into FlickR training is provided by Figure 9, which illustrates the output of the FlickR trained model, improvement with infusion of 5% camera trap images, and the output of alternative camera trap dataset training (South Africa, Tanzania, North America and Europe). Without the addition of camera trap images, the FlickR model achieved AP results of 68.06% on the Australian dataset, detecting all but the most difficult examples in the test set. With 5% infusion (30 trap images), the mAP increased by 12.47% to 80.53%. In contrast, the camera trap model trained on images from camera traps in North America, Europe, South Africa and Tanzania only achieved an AP of 68.96% on the Australian test set. These results suggest that FlickR training with the addition of trap images in cases of underperformance is a more appropriate method of training DCNNs for camera trap image processing than camera trap training alone.

## 4. Discussion

We investigated the use of non-iconic FlickR images as an alternative to annotated camera trap images in the task of DCNN training for camera trap image processing tasks. Specifically, we established the greater transferability of the FlickR trained model when compared to models trained on single location data, and the competitiveness of FlickR training in comparison to training on a variety of camera trap images. We then demonstrated how such a model can be optimized for specific camera trap applications, with minimal annotation of camera trap data, to achieve optimal results via infusion of camera trap images into FlickR training. Our results show that FlickR training significantly improves model robustness and location invariance. In particular, it provides ecologists with a practical, cost effective, out of the box solution, capable of detecting pigs even in the most challenging camera trap environments. We established the effectiveness of camera trap image infusion into the FlickR trained model to further improve performance. This suggests that ecologists can deploy camera traps to collect data, run the FlickR trained model on collected data, and if it is necessary, optimize the model using their own data. An interesting area of research would be integration of this functionality within ClassifyMe (Falzon, Lawson et al. 2020) which would allow researchers to download a FlickR model, apply it to their camera trap data on their PC, and in the case of underwhelming performance, submit a small dataset of their images for optimization training. In light of the growing number of camera trap based projects undertaken by ecologists, this research provides an invaluable method by which researchers can process extensive image data regardless of their location, camera trap environment or the particularities of the subspecies they are studying.

One problem commonly faced by ecologists is the lack of camera trap images of rare species (Willi, Pitman et al. 2018). This poses significant problems when training multi-class object detectors, as the large class imbalance between common species (many images) and rare species (very few images) causes object detectors to misclassify species, over enthusiastically classifying species based on how common they were on the dataset. This was observed by (Willi, Pitman et al. 2018), who noted that strong class imbalance causes the DCNN to misclassify species by favouring very common classes. For example, they noted that insufficient images of the rare striped hyena resulted in their model achieving an AP of 0% on this class. This limitation in data can be rectified using images of rare species sourced from FlickR. A search of the term ‘striped hyena’ on FlickR returns over 2,700 images which could have been used in training their model to rectify the class imbalance. It is worth noting however that not all search results are accurately labelled which means the FlickR images would have to be sorted prior to training. This may be an interesting area of future research.

The use of FlickR images as the principal training data also rectifies another major problem faced by researchers. Studies have indicated that DCNNs have a tendency to return overly confident predictions (Willi, Pitman et al. 2018). We observed this phenomenon when models were trained on camera trap data alone. This may be due to the high consistency in image quality, lighting, camera angle and vegetation in camera trap data, particularly if the only foreground object/s are pigs. Furthermore, many trap images feature obscured or poor-quality imagery of pigs which if used in the training set, may cause the network to make unrealistically optimistic predictions, by attributing 100% confidence to visual features which may not display sufficiently distinct characteristics present solely in the *Suidae* class. In contrast, the higher resolution of FlickR images and large variations between images forces the model to reduce the confidence attributed to poor quality or obscured pigs.

It is noteworthy that to the best of our knowledge, researchers in this field do not use explicit negative sampling in the training of DCNNs for camera trap image processing. Although the majority of studies focus on multi-class training, which reduces the need for negative sampling due to implicit inter-class discrimination, no studies have examined the impact of inclusion or exclusion of explicit negative sampling. Lack of negative sampling can cause DCNNs to over-enthusiastically classify objects, without sufficient ability to discriminate against out of sample species. For example, if a network is trained to classify wildebeest, pigs and elephants, it may exhibit a tendency to classify rhinoceros as belonging to either of the three learned classes. If negative samples of rhinoceros were included, this would minimise misclassification. This issue may be a fruitful area of research.

This research did not investigate the application of the FlickR training method using alternative object detectors such as YOLOv3 (Redmon and Farhadi 2016), and Faster R-CNN (Ren, He et al. 2015). RetinaNet was chosen as it achieves a balance between the computational efficiency of YOLOv3 and the accuracy of Faster-RCNN, which made it an appropriate choice for the difficult task of camera trap image processing.

The solution provided by this study could be improved by exploring object segmentation. Object segmentation builds upon the benefits of object detection by largely eliminating background influence on model performance, thus improving model accuracy and overall performance. Nevertheless, this research is an important improvement over most camera trap solutions which achieve image classification rather than object detection.

## 5. Conclusion

This study investigated the useability of animal images obtained from the consumer image sharing website FlickR in the task of training Deep Convolutional Neural Networks for camera trap image processing. It established the benefits of using FlickR images in improving location invariance and robustness to environmental features. Furthermore, it evaluated the use of camera trap image infusion into FlickR datasets for DCNN training. This research provides ecologists with a data collection and model training method which could drastically reduce the data collection, annotation and model training costs associated with processing camera trap imagery. In effect, this contribution provides a means by which Artificial Intelligence can be deployed in ecological management on a large scale.

## Supporting information

Muliticlass Application

SSIM Duplicate remover pseudocode

Training Parameters Transfer Learning

Negative Sampling

Review of Classification studies

## 6. Acknowledgements

Andrew Shepley is supported by an Australian Postgraduate Award. We would like to thank the Australian Department of Agriculture and Water Resources, the Centre for Invasive Animals Solutions, University of New England and the NSW Department of Primary Industries for supporting this project. We appreciate the Creative Commons Images provided through FlickR; Australian camera trap images provided to us by Mark Lamb and Jason Wishart; the Snapshot Serengeti, University of Missouri Camera Traps and North American Camera Traps datasets through the Labeled Information Library of Alexandria: Biology and Conservation and the Camera CATalogue dataset provided through the Data Repository of the University of Minnesota.

## 7. Author Contributions

Andrew Shepley, Greg Falzon and Paul Kwan conceived the ideas and designed methodology; Andrew Shepley, Paul Meek and Greg Falzon collected the data; Andrew Shepley and Greg Falzon analysed the data; Andrew Shepley led the writing of the manuscript. All authors contributed critically to the drafts and gave final approval for publication.

