## Supplementary material for "Location Invariant Animal Recognition Using Mixed Source Datasets and Deep Learning": Muliticlass Application

### APPENDIX S5

##### Application to Real Life Multi-Class Detection Problem

To demonstrate the application of this training method to multi-class tasks, we trained a Retinanet model according to the specifications provided in Appendix S4. We downloaded 300 images of each of the classes pig, kangaroo, fox and goat from FlickR, along with approx. 1000 negative samples. Search queries for both positive and negative samples are provided in Table 1.

| *Positive Search Queries* | *Negative Search Queries* |
|---|---|
| *Kangaroo* | *Dingo* |
| *Kangaroos* | *Cat* |
| *Eastern Grey Kangaroo* | *Chat sauvage* |
| *Macropus giganteus* | *Raccoon* |
| *Infrared kangaroo* | *Antelope* |
| *Feral goat* | *Buffalo* |
| *Feral goats* | *Rabbits* |
| *Capra hircus* | *Rhinoceros* |
| *Fox* | *Grizzly bear* |
| *Foxes* | *Bison* |
| *Vulpes Vulpes* | *Wildebeest* |
| *Infrared fox* | *Rats* |
| *Australian fox* | *Lizards* |
| For search queries for class 'Pig' refer to Section 2.1(i) Table 1 | *Deer* |
| | *Birds* |
| | *Crows* |
| | *Chimpanzees* |
| | *Stones* |

**Table 1:** *FlickR search queries for positive and negative samples*

The model was then tested on an out of sample trap site dataset (Scotts) with class distribution illustrated by Figure 1. Note that classes with less than 10 instances were ignored for practical purposes.

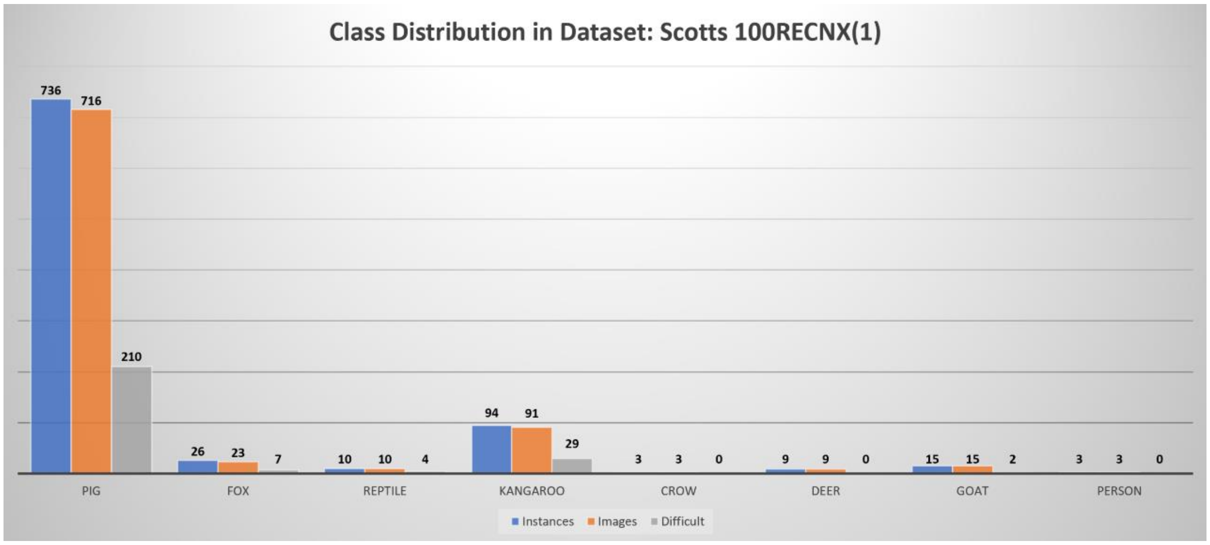


***Figure 1:*** *Class distribution of out of sample Scotts multi-class test set. The class pig was the most commonly observed class. All classes with 10 or less instances were ignored.*

Training solely on FlickR images resulted in a sub-optimal mAP as illustrated by Table 2.

| CLASS | AP (%) |
|---|---|
| ***Pig*** | 90.87 |
| ***Kangaroo*** | 66.83 |
| ***Goat*** | 66.34 |
| ***Fox*** | 23.76 |
| **mAP (%)** | 61.95 |

**Table 1:** *Results of multi-class training without infusion*

As such, we applied the camera trap infusion training method outlined in Section 2.2 (ii). Infusion images were obtained from camera trap sites in Kilparney and Yarra. Only 30 images from each class were required (10% infusion) to achieve an 18.99% increase in mAP on the out of sample Scotts dataset. The fox class mAP improved the most, with an increase of 42.76%. The mAP improved across all classes, as shown by Table 2.

| CLASS | AP (%) |
|---|---|
| ***Pig*** | 94.61 |
| ***Kangaroo*** | 76.85 |
| ***Goat*** | 85.77 |
| ***Fox*** | 66.52 |
| **mAP (%)** | 80.94 |

**Table 2:** *Results of multi-class training with 10% infusion*

Sample output images are provided in Figures 2 and 3.

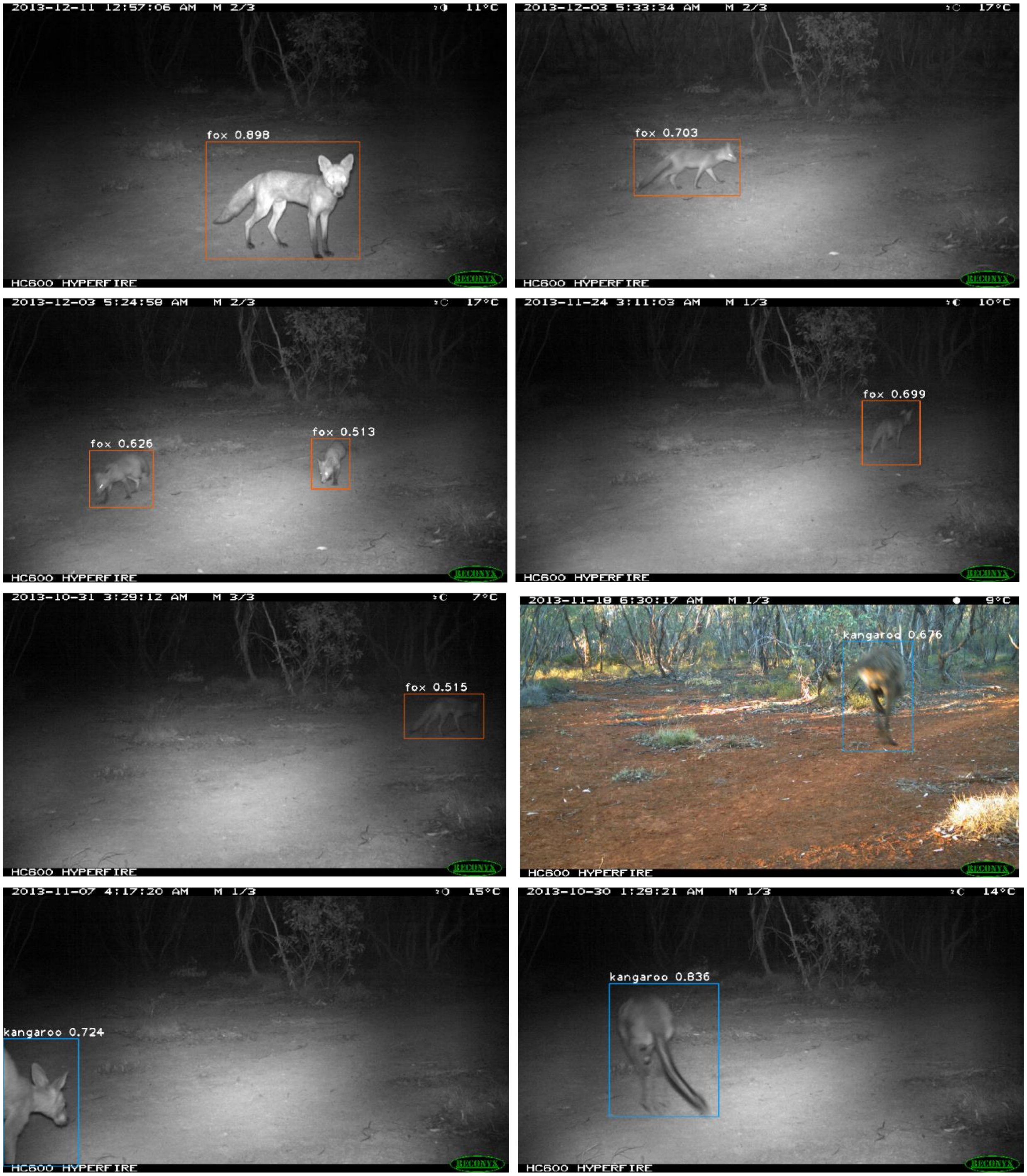

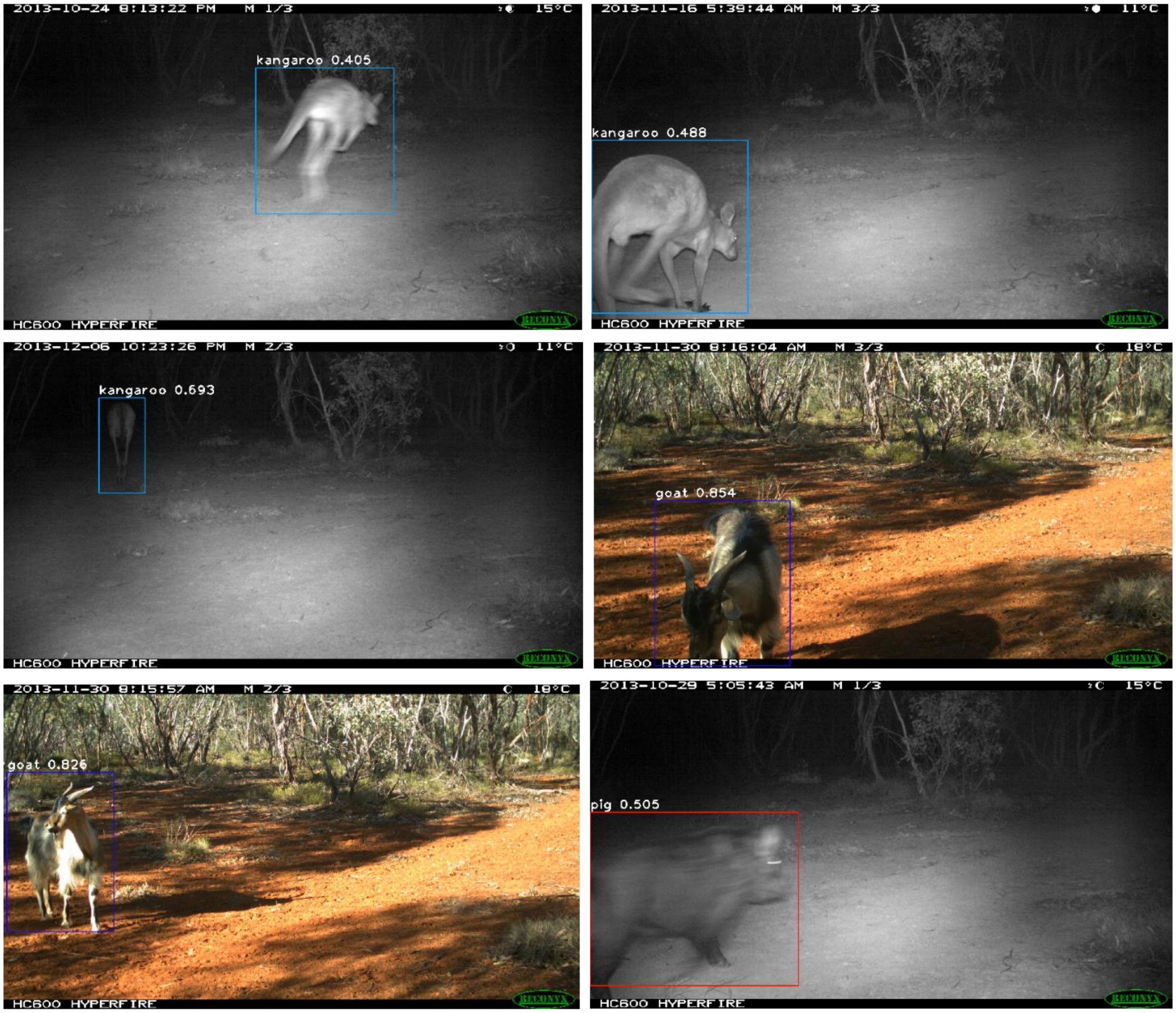
2013-10-24 8:13:22 PM M 1/3 15°C
kangaroo 0.405
HC600 HYPERFIRE
2013-11-16 5:39:44 AM M 3/3 11°C
kangaroo 0.488
HC600 HYPERFIRE
2013-12-06 10:23:26 PM M 2/3 11°C
kangaroo 0.693
HC600 HYPERFIRE
2013-11-30 8:16:04 AM M 3/3 18°C
goat 0.854
HC600 HYPERFIRE
2013-11-30 8:15:57 AM M 2/3 18°C
goat 0.826
HC600 HYPERFIRE
2013-10-29 5:05:43 AM M 1/3 15°C
pig 0.505
HC600 HYPERFIRE

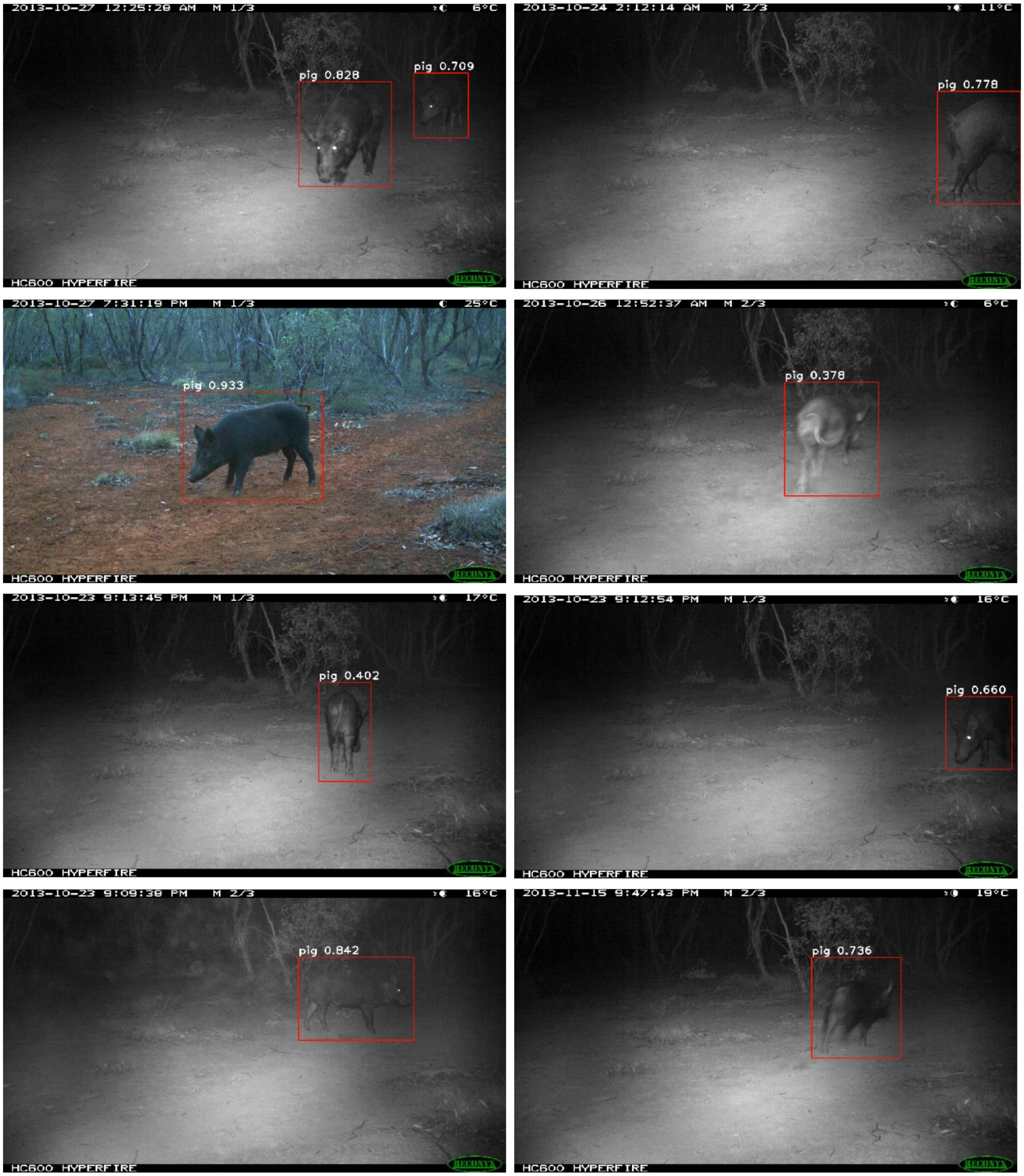


**Figure 2:** *Correctly classified and located classes*

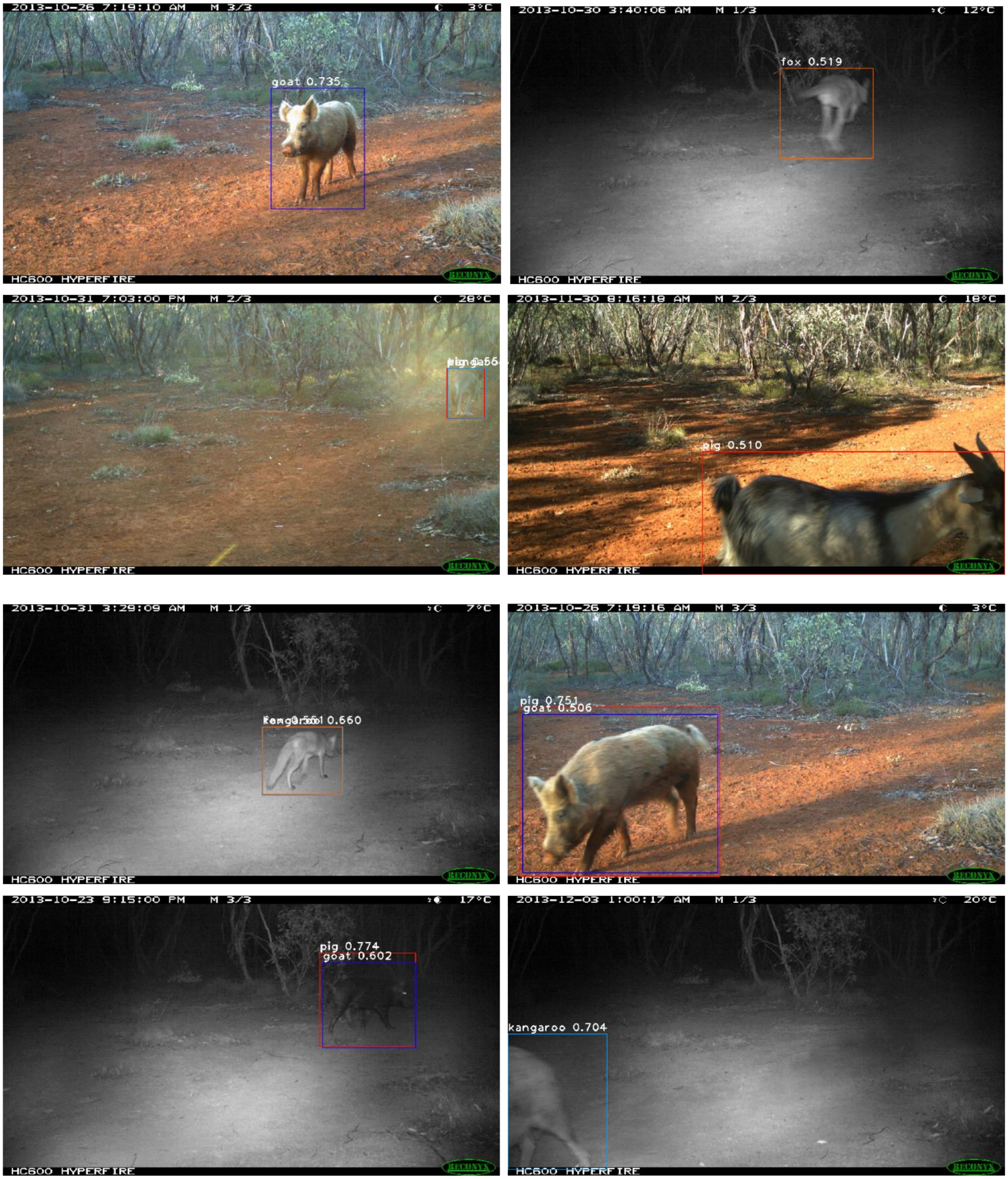


**Figure 3:** *Incorrectly classified classes.*
