## Supplementary material for "Location Invariant Animal Recognition Using Mixed Source Datasets and Deep Learning": SSIM Duplicate remover pseudocode

### APPENDIX S4: SSIM Image Duplicate Remover

Andrew Shepley

#### I. Algorithm Pseudocode

**Algorithm 1** SSIM - Duplicate Remover

**Input:** $dir1$, $dir2$, $dir3$
$dir1$ is the directory containing all images
$dir2$ is an empty directory to contain unique images
$dir3$ is an empty directory to contain duplicate images
**begin:**
**for** $f_i$ in $dir1$ **do**
  $imageA \leftarrow f_i$
  **while** $dir1$ contains more than one file **do**
    **if** $Algorithm2(dir1, imageA)$ is $True$ **then**
      $dir3 \leftarrow dir3 \cup f_i$
    **else**
      $dir2 \leftarrow dir2 \cup f_i$
    **end if**
    $dir1 \leftarrow dir1 - f_i$
  **end while**
  **if** $length(dir1) == 1$ **then**
    **if** $Algorithm2(dir2, imageA)$ is $True$ **then**
      $dir3 \leftarrow dir3 \cup f_i$
    **else**
      $dir2 \leftarrow dir2 \cup f_i$
    **end if**
    $dir1 \leftarrow dir1 - f_i$
  **end if**
**end for**
**return** $dir2$

**Algorithm 2** SSIM - Duplicate Remover

**Input:** $dir$, $imageA$
$dir1$ is the directory containing all images
$image$ is an image to compare to all images in the directory
**begin:**
**for** $f_i$ in $dir$ **do**
  **if** $f_i$ != $imageA$ **then**
    $imageB \leftarrow f_i$
    $score \leftarrow compareSSIM(imageA, imageB)$
    **if** $score == 1$ **then**
      **return** $True$
    **end if**
  **end if**
**end for**
**return** $False$
