## Supplementary material for "Location Invariant Animal Recognition Using Mixed Source Datasets and Deep Learning": Training Parameters Transfer Learning

### APPENDIX S3

#### Model Architecture and Training Parameters

The majority of ecological deployments of deep learning classifiers involve training of a ResNet model on a custom dataset (Norouzzadeh, Nguyen et al. 2017, Willi, Pitman et al. 2018, Tabak, Norouzzadeh et al. 2019). We trained a RetinaNet model (Lin, Goyal et al. 2018) with a ResNet-50 backbone using standard object detection training parameters. RetinaNet is a state-of-the-art object detector, commonly used in general object detection tasks (see Figure 1), and in specific applications such as face detection and recognition (Wang, Yuan et al. 2017), pedestrian detection (Milton 2018), and autonomous vehicles (Hoang, Nguyen et al. 2019). It boasts accuracy similar to computationally expensive 2 stage object detectors such as Faster R-CNN, with the benefit of speed similar to Single Shot Detectors (SSD) (Chowdhury, Garg et al. 2019).

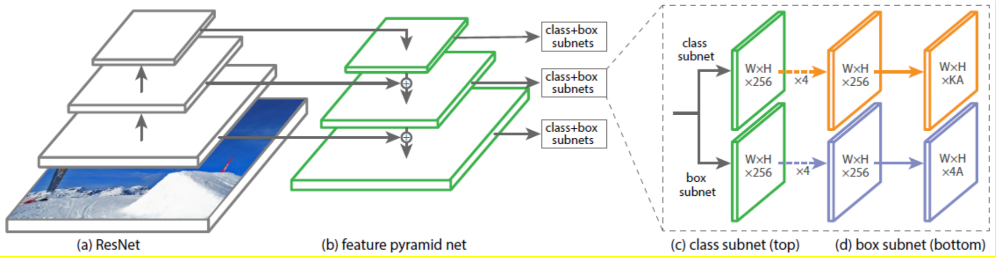


**Figure 1**: *RetinaNet architecture: A Feature Pyramid Network (FPN) on top of a feedforward ResNet architecture. Two classification subnetworks achieve bounding box regression and object classification (Lin, Goyal et al. 2018)*

The experimental results provided by this research were collected using the Keras implementation of RetinaNet (with default ResNet-50 backbone) provided by the authors of RetinaNet, including a focal loss for classification and detection model pretrained on the MS COCO dataset (Zhang, Wen et al. 2018). This model was employed to achieve transfer learning using the Tensorflow graph computation framework (Abadi, Barham et al. 2016). We used a batch size of 8, 500 steps, and 50 epochs (Lin, Maire et al. 2014). Random transform was applied, and the backbone was frozen. All training and testing was conducted using a Centos7.7 Linux virtual machine, with 16 x 2.4GHz Intel Xeon processors, 128GB memory, and Tesla V100-PCIE-32GB GPU.

#### Transfer Learning

Rather than training our model from scratch on a large ecological dataset such as Snapshot Serengeti, we repurposed general features learned from the MS COCO dataset (Lin, Maire et al. 2014) to the task of pig detection. The MS COCO dataset contains 328k images including 2.5 million labelled instances of 80 classes. This includes classes such as ‘sheep’, ‘cow’, ‘horse’, and ‘dog’ which share features similar to those of pigs. According to (Yosinski, Clune et al. 2014), transfer learning with backbone freezing is the most appropriate form of training when using a small dataset with high resemblance to classes learned by the pretrained network. This avoids overfitting, by preserving learned features, while training the classification subnet to reclassify learned features based on their similarity with features of the pig class.

This study provides results of transfer learning experiments on a ResNet-50 backbone only, as there were no publicly available pretrained ResNet-101 backbones for RetinaNet, which meant training from scratch would be required. This is a very computationally expensive and data intensive process, requiring tuning of learning parameters to achieve optimal results (Hoang, Nguyen et al. 2019). We considered this was unnecessary as transfer learning achieved excellent results.
