## Supplementary material for "Location Invariant Animal Recognition Using Mixed Source Datasets and Deep Learning": Negative Sampling

### APPENDIX S2: Negative Sampling

The presence of negative samples in the training dataset is an essential component of any training pipeline (Ren, He et al. 2015). This is particularly important in single-class object detection (Gao, He et al. 2019) as it prevents over-enthusiastic classification of objects by allowing the network to discriminate between positive and negative samples. During multi-class object training, class characteristics are automatically differentiated during training. In contrast, the absence of alternative classes in single class object detection encourages the network to classify indiscriminately.

Thus, it is necessary to train DCNNs to recognise the distinctive features of pigs by training it to recognise which features are not attributable to pigs. For example, wildebeest and bison share very similar features, and inhabit similar ecosystems to warthogs. Therefore, it is extremely important to ensure a significant set of negative samples containing objects with similar characteristics to the chosen class is used during training. To rectify this problem, we downloaded 878 images of 35 non-pig animal species, including humans, as illustrated by Table 1. These images were included in all training sets for all single class trained models in this study.

| Species | Nº images | Species | Nº images |
|---|---|---|---|
| 1. Chimpanzee | 30 | 20. Elephant | 14 |
| 2. Meerkat | 31 | 21. Buffalo | 17 |
| 3. Antelope | 32 | 22. Wildebeest | 15 |
| 4. Lion | 16 | 23. Rat | 13 |
| 5. Hippopotamus | 30 | 24. Stumps | 16 |
| 6. Rhinoceros | 30 | 25. Stones | 15 |
| 7. Chipmunk | 30 | 26. Car | 16 |
| 8. Goat | 33 | 27. Bird | 16 |
| 9. Rabbit | 15 | 28. Turtle | 14 |
| 10. Bison | 35 | 29. Cat | 14 |
| 11. Grizzly Bear | 36 | 30. Dingo | 15 |
| 12. Kangaroo | 33 | 31. Dog | 16 |
| 13. Wallaby | 33 | 32. Sheep | 16 |
| 14. Boulders | 9 | 33. Horse | 15 |
| 15. Raccoon | 16 | 34. Bear | 30 |
| 16. Tiger | 14 | 35. Giraffe | 16 |
| 17. Zebra | 16 | 36. Gorilla | 16 |
| 18. Human | 119 | 37. Polar Bear | 14 |
| 19. Moose | 16 | 38. Hare | 16 |
| **878 negative samples** | | | |

**Table 1**: *Number of negative samples according to species type*

The use of negative sampling incidentally solved another problem faced by ecologists. Differentiating between empty frames, and those containing a species of interest is often approached as a separate task to object detection. For example, (Willi, Pitman et al. 2018) trained two separate models; one was used to sort empty frames from those containing species of interest, and another to classify species. This resulted in a positive bias in the species trained model, which led to misclassification. In contrast, we trained our models to automatically differentiate empty frames via inclusion of explicit negative sampling in the training set.

During preliminary experimentation, we found that due to the limited presence of non-pig species within the FlickR and camera trap datasets, the trained models over enthusiastically classified any animal-like or foreground object as belonging to the class ‘pig’. Our experimental results strongly indicate that without a negative sample dataset containing images of bears and dogs, the network will classify instances of these species as ‘pig’ with relatively high confidence, as shown in Figure 1.

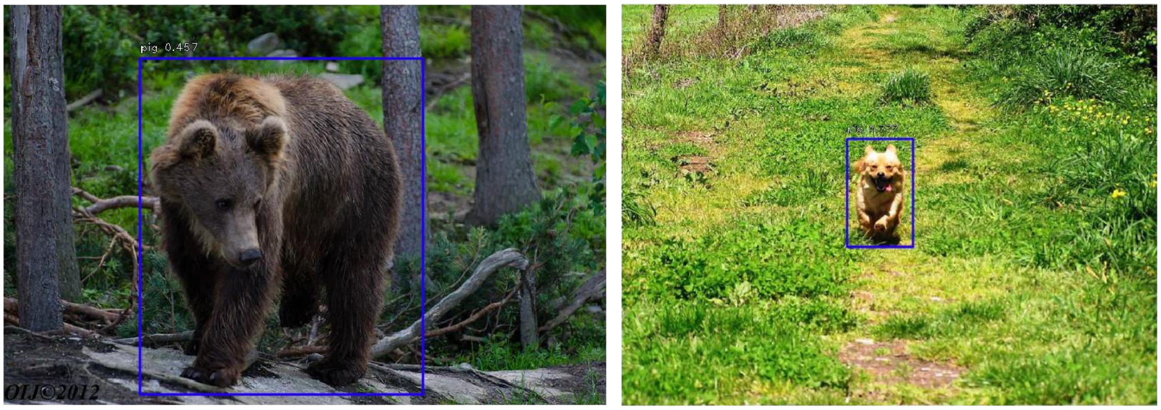


**Figure 1:** *Without explicit negative sampling, the network classified the bear on the left as a pig, with 45.7% confidence. Similarly it classified the dog on the right as a pig, with 77.7% confidence.*

Experimental results suggest that the ratio of negative samples to positive samples should not be below 2:1. An interesting area of future research would be to determine the optimum ratio of negative to positive samples in single class object detection. Another potential area of research would be to examine whether explicit negative sampling is beneficial in multi-class object detection.
