## Supplementary material for "Location Invariant Animal Recognition Using Mixed Source Datasets and Deep Learning": Review of Classification studies

### APPENDIX S1: Review of Image Classification Solutions

(Gomez Villa, Salazar, & Vargas-Bonilla, 2016) proposed a method to automate species identification in a simplified version of the Snapshot Serengeti dataset, aiming to reduce the time and resource expenditure dedicated to analysis of large volumes of camera trap images. They experimented on six state of the art DCNNs, achieving accuracies ranging from 35.4% to 88.9% accuracy in the Top-1 classification task. Higher accuracies were only achieved when a balanced dataset containing only manually segmented foreground animals was used. This work was built upon by (Norouzzadeh et al., 2017) which used transfer learning to address factors such as the number of species in a given image, the presence of young, and animal behaviour. They achieved superior results (93.8%) on the Snapshot Serengeti dataset, but did not report results on out of sample data, meaning the robustness of the model was not evaluated. The transferability of the model development procedure is also limited by dependency on large datasets, which is not realistic in most practical ecology applications. Similarly, (Nguyen et al., 2017) performed species classification, comparing the performance of transfer learning and training from scratch. They used a subset of 107,022 images of 15 species from the Wildlife Spotter dataset (www.ala.org.au). They achieved AP results ranging from 84.39-96%. However, like (Norouzzadeh et al., 2017), they did not evaluate their model on an out of sample set, therefore its robustness cannot be assessed.

(Willi et al., 2018) attempted to address this issue by investigating the use of transfer learning to classify images from smaller camera trap datasets. They trained a model from scratch using the Snapshot Serengeti dataset, and then used transfer learning to repurpose the learned features for the task of image classification on the smaller trap datasets. They demonstrated that transfer learning does result in improved accuracy, however, this process is limited in value, as their model could not generalise well to other camera trap datasets. The model may have learned specific features and species biases within their training set. They noted that their model could only be used to classify images from the traps used for training and would need to be retrained for use in other camera traps characterised by differences in vegetation, camera placement, study sites and species. This limitation in model robustness and invariance to location and species variations is a significant factor inhibiting the deployment of DCNN models in camera trap applications. To address this, we propose the use of FlickR images to provide variety in background and image context to broaden the useability of trained models, thus shifting emphasis from quantity of data to quality of data. Furthermore, both (Norouzzadeh et al., 2017) and (Willi et al., 2018) developed deep learning solutions aimed for use alongside citizen science. In contrast, this study proposes a fully automated deep learning system, without reliance on citizen science.

To develop an image classification solution that could be used across all camera trap sites, (Tabak et al., 2019) trained a ResNet model on 3,741,656 images including 27 species from 5 locations in the US. They achieved 97.6% accuracy on test images from these sites. However, despite using data augmentation techniques such as cropping, flipping, translation and brightness changes, their model only achieve 82% on out of sample test images. They acknowledged that the model was not location or background invariant and proposed training on all possible environments would be needed to increase accuracy. In an attempt to identify the underlying cause of location invariance (Miao et al., 2019) used gradient-weighted class-activation mapping (Grad-CAM) to illustrate the most salient features used by the network to classify species. This investigation provided evidence that DCNNs learn to associate the presence of environmental factors, such as trees, with particular species if they appear frequently in the training dataset. This inner dataset bias degrades the ability of the network to generalise to other datasets.
